# Ambient-Condition Visualization of Metalloenzyme Reactive Intermediates Enabled by the SHINERS Paradigm

**DOI:** 10.64898/2026.10.03.756395

**Authors:** Langxing Liao, Qiong-Qiong Li, Xuan Zhang, Jia Liu, Zi-Han Chen, Yang Zhou, Zongyu Bao, Hao Ma, Liulin Yang, Xiang Wang, Cheng Wang, Bin Ren, Jian-Feng Li, Binju Wang

**Affiliations:** State Key Laboratory of Physical Chemistry of Solid Surfaces, College of Chemistry and Chemical Engineering, Fujian Provincial Key Laboratory of Theoretical and Computational Chemistry, Collaborative Innovation Center of Chemistry for Energy Materials (iChEM), College of Energy, College of Materials, College of Life Science, Institute of Artificial Intelligence, Xiamen University, Xiamen 361005, China; School of Pharmaceutical Sciences, Hainan University, Haikou 570228, China; College of Chemistry and Materials Science, Fujian Normal University, Fuzhou 350108, China; Innovation Laboratory for Sciences and Technologies of Energy Materials of Fujian Province (IKKEM), Xiamen University, Xiamen 361005, China

**Author notes:** **Corresponding author.** (B.W.); (J.-F.L.). These authors contributed equally to this work (Langxing Liao, Qiong-Qiong Li, Xuan Zhang, Jia Liu).

## Abstract

Reactive metal–oxygen intermediates are central to metalloenzyme catalysis, yet their low populations and short lifetimes make vibrational characterization under aqueous reaction conditions challenging. Here, we establish shell-isolated nanoparticle-enhanced Raman spectroscopy (SHINERS) as a powerful vibrational approach for visualizing reaction-associated metal–ligand signatures of reactive metalloenzyme states. Across the heme and non-heme iron enzymes TauD, VioC, CYP119, and TqaL_Nc_, isotope-sensitive features provide vibrational information on reactive coordination states. In TauD and VioC, isotope-sensitive features provide vibrational characterization of Fe(IV)=O intermediates. In CYP119, an isotope-sensitive Fe(IV)=O vibration is assigned to P450 Compound I generated by *m*-CPBA, including under lauric-acid turnover conditions. Most notably, in TqaL_Nc_, combined ^18^O- and ^15^N-sensitive vibrational responses support formation of the proposed (R)HN–Fe(IV)=O coordination state and the dynamic-coordination mechanism underlying selective C–N bond formation. These results demonstrate that SHINERS can serve as a powerful technique for the in situ vibrational visualization of reactive intermediates and their coordination states in metalloenzyme catalysis.

---

Reactive oxygen intermediates generated by metalloenzymes play central roles in biological oxidation chemistry, enabling challenging transformations including C–H hydroxylation, desaturation, halogenation, and C–N/C–C bond formation^1–6^. Iron-dependent heme and non-heme enzymes (Fig. 1a) are particularly prominent owing to their ability to activate molecular oxygen and generate highly reactive metal–oxygen species such as Fe(III)–OO^−^, Fe(IV)=O, and porphyrin radical cation intermediates^3–12^. These transient species are widely recognized as the key catalytic oxidants in enzymatic oxidative transformations.

**Fig. 1.**
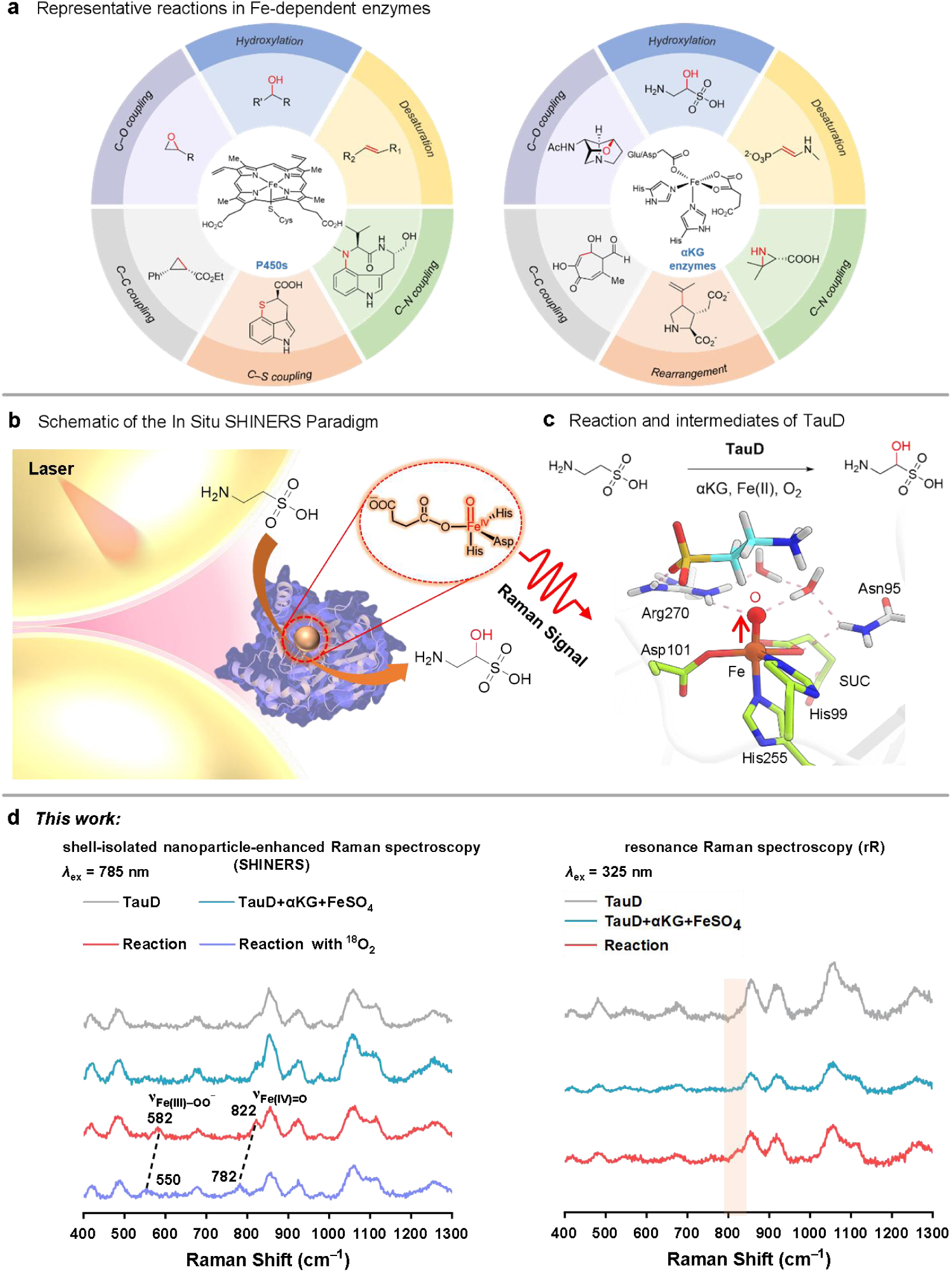
In situ SHINERS characterization of reactive metal–oxygen species in metalloenzyme catalysis. **a**, Representative reactions catalyzed by Fe-dependent P450s and αKG-dependent non-heme enzymes. **b**, Schematic of the interfacial SHINERS configuration, in which Au@SiO_2_ shell-isolated nanoparticles enhance Raman scattering from molecular species positioned within the plasmonic near-field, including interfacially accessible enzyme states. **c**, TauD-catalyzed reaction and QM/MM-optimized structure of the Fe(IV)=O intermediate. **d**, SHINERS spectra of the TauD reaction (left; *λ*_ex_ = 785 nm), showing an oxygen-isotope-sensitive lower-frequency feature consistent with an Fe(III)–peroxo state and a higher-frequency feature assigned to Fe(IV)=O, together with resonance Raman spectra collected without SHINs for comparison (right; *λ*_ex_ = 325 nm). For the oxygen-isotope measurement, the TauD reaction was performed under an ^18^O_2_ atmosphere under otherwise matched SHINERS conditions. SHINERS conditions: 200 μM TauD, 1 mM taurine, 2 mM αKG, 200 μM FeSO4, 150 mM Tris-HCl (pH 8.0), 300 mM NaCl, 4 °C. Resonance Raman conditions: 400 μM TauD, 1 mM taurine, 2 mM αKG, 400 μM FeSO4, 150 mM Tris-HCl (pH 8.0), 300 mM NaCl, −40 °C.

A broad range of spectroscopic, structural, and single-molecule methods has provided fundamental insight into reactive metalloenzyme intermediates. Stopped-flow absorption spectroscopy defines electronic and kinetic changes after rapid mixing; time-resolved resonance Raman spectroscopy provides mode-specific vibrational information; Mössbauer and EPR/ENDOR methods report on oxidation state, spin state, and electronic structure; X-ray absorption and emission spectroscopies probe metal-site geometry and electronic structure; serial crystallographic and XFEL approaches can resolve structural states; and single-molecule conductance measurements can distinguish individual catalytic states through characteristic electrical signatures^13–33^. These methods probe distinct observables and temporal regimes, making convergent evidence particularly important for short-lived or low-population enzymatic states.

Raman spectroscopy directly reports molecular vibrations, and resonance enhancement has been especially valuable for chromophoric heme intermediates^34,35^. Surface-enhanced Raman spectroscopy (SERS) can increase scattering from molecules located within a plasmonic near-field^36–38^. SHINERS extends this concept by coating plasmonic nanoparticles with an ultrathin, chemically isolating shell, thereby reducing direct contact between the metal core and the sample while retaining near-field electromagnetic enhancement^39–47^. This architecture enables vibrational interrogation at solid–liquid interfaces and access to weak metal–ligand signatures in aqueous catalytic systems. Although SHINERS is well established in surface and interfacial catalysis, its application to reactive metalloenzyme states remains largely unexplored. These features make SHINERS well suited for probing metal–ligand vibrations associated with enzymatic states under aqueous reaction conditions.

Here, we use in situ SHINERS to access the metal–ligand vibrational signatures of reactive intermediates in representative heme and non-heme iron enzymes under aqueous catalytic conditions. Vibrational assignments are established through isotope substitution, reaction and background controls, independent spectroscopy where applicable, and harmonic QM/MM normal-mode calculations. Across TauD and VioC, SHINERS resolves isotope-sensitive vibrational signatures of Fe(IV)=O intermediates, while in CYP119 an isotope-sensitive Fe(IV)=O vibration is assigned to P450 Compound I. The principal mechanistic advance emerges from TqaL_Nc_, where combined oxygen- and nitrogen-isotope responses support the proposed (R)HN–Fe(IV)=O state and provide an experimental anchor for the dynamic-coordination mechanism underlying selective C– N bond formation. Overall, these results establish SHINERS as a vibrational strategy for probing reactive intermediates and their coordination states in metalloenzyme catalysis.

## Results

### In situ SHINERS accesses reactive metal–ligand vibrations in aqueous metalloenzyme reactions

Fig. 1b illustrates the interfacial in situ SHINERS configuration used in this study. Au@SiO_2_ shell-isolated nanoparticles (SHINs), consisting of an Au core surrounded by an ultrathin silica shell, were characterized by SEM and TEM (Supplementary Fig. 1) and UV–visible extinction spectroscopy (Supplementary Fig. 2). For Raman measurements, the SHINs were deposited on a glassy-carbon support and brought into contact with the enzyme-containing aqueous phase, while a confocal Raman microscope focused the excitation and collection volume on the SHIN-rich solid/liquid interface (Supplementary Fig. 3). Here, “in situ” refers to spectral acquisition from the reacting enzyme mixture at the SHIN-coated interface. The ultrathin silica shell isolates the Au core from direct contact with the biomolecular sample while allowing the plasmonically enhanced electromagnetic near-field to extend into the aqueous phase, providing an interfacial configuration for Raman interrogation of enzyme-associated vibrational states^40,42,46^.

We first benchmarked this configuration using taurine/α-ketoglutarate (αKG) dioxygenase (TauD), a Fe(II)/αKG-dependent oxygenase with well-characterized high-valent iron–oxygen intermediates^48–50^. TauD retained catalytic activity in the presence of SHINs, with no substantial loss of product formation observed in activity assays (Supplementary Figs. 5–7). Reference spectra of the individual small-molecule reaction components were collected to define the spectral background (Supplementary Fig. 8), while spectra of TauD alone and the pre-reaction TauD/αKG/Fe^2+^ mixture were obtained as reference states for subsequent analysis (Fig. 1d).

Following addition of taurine, two reaction-associated features appeared near 582 and 822 cm^−1^ (Fig. 1d, left). Under an ^18^O_2_ atmosphere, these features shifted to approximately 550 and 782 cm^−1^, respectively. The approximately 40 cm^−1^ downshift of the higher-frequency band is close to the approximately 36 cm^−1^ value expected from the reduced-mass relation for an Fe–O vibration (Supplementary Table 1; Methods), while harmonic QM/MM calculations yielded an Fe–O stretching mode near 825 cm^−1^ for the Fe(IV)=O intermediate (Supplementary Fig. 9). The lower-frequency feature also exhibited an oxygen-isotope response and falls within the spectral region previously associated with Fe(III)–peroxo species^51^. Collectively, the reaction dependence, oxygen-isotope responses and QM/MM analysis establish the approximately 822 cm^−1^ band as an Fe(IV)=O vibration and support assignment of the near 582 cm^−1^ feature to an Fe(III)–peroxo-associated mode.

Notably, SHINERS enhancement is not intrinsically selective for ferryl vibrations over other protein modes, and its enhancement depends on interfacial geometry, molecular orientation, and mode-specific Raman polarizability. Accordingly, assignment of the metal–oxygen vibrations was based on the combined evidence from reaction-dependent spectral changes, reproducible oxygen-isotope shifts, background controls and agreement with QM/MM normal-mode calculations. For instance, bands near approximately 490 and 840 cm^−1^ were also present in the enzyme-background spectra but showed no resolved oxygen-isotope-dependent frequency shifts, distinguishing them from the isotope-sensitive Fe–O features associated with the reactive intermediates.

For comparison, conventional resonance Raman spectra were collected without SHINs using 325 nm excitation under the stated experimental conditions (Fig. 1d, right). Under these measurements, a comparably distinct feature near 822 cm^−1^ was not resolved, whereas SHINERS provided direct access to the isotope-responsive Fe–O vibrations in the aqueous TauD reaction. The importance of the silica shell was further examined using uncoated Au nanoparticles, which produced substantial changes in the overall Raman background and did not yield a reproducible ferryl-associated feature (Supplementary Fig. 11). These benchmark experiments establish a SHINERS configuration capable of resolving reaction-associated and isotope-sensitive metal–oxygen vibrations in an enzymatic system under aqueous conditions.

### Extension of in situ SHINERS to heme and non-heme iron enzyme systems

We next extended the SHINERS analysis to the Fe(II)/αKG-dependent hydroxylase VioC (Fig. 2a) and the thermophilic cytochrome P450 CYP119 (Fig. 2d), representing non-heme and heme iron systems, respectively. VioC is proposed to employ a ferryl Fe(IV)=O intermediate during L-arginine hydroxylation^52^, whereas reaction of CYP119 with *m*-chloroperoxybenzoic acid (*m*-CPBA) provides a well-established route to highly reactive P450 Compound I chemistry^14,53^. Although Fe(IV)=O stretching vibrations of Compound I have been identified by resonance Raman spectroscopy in related heme enzymes such as peroxidases^54^, direct vibrational characterization of the Fe(IV)=O unit in P450 Compound I has remained elusive.

**Fig. 2.**
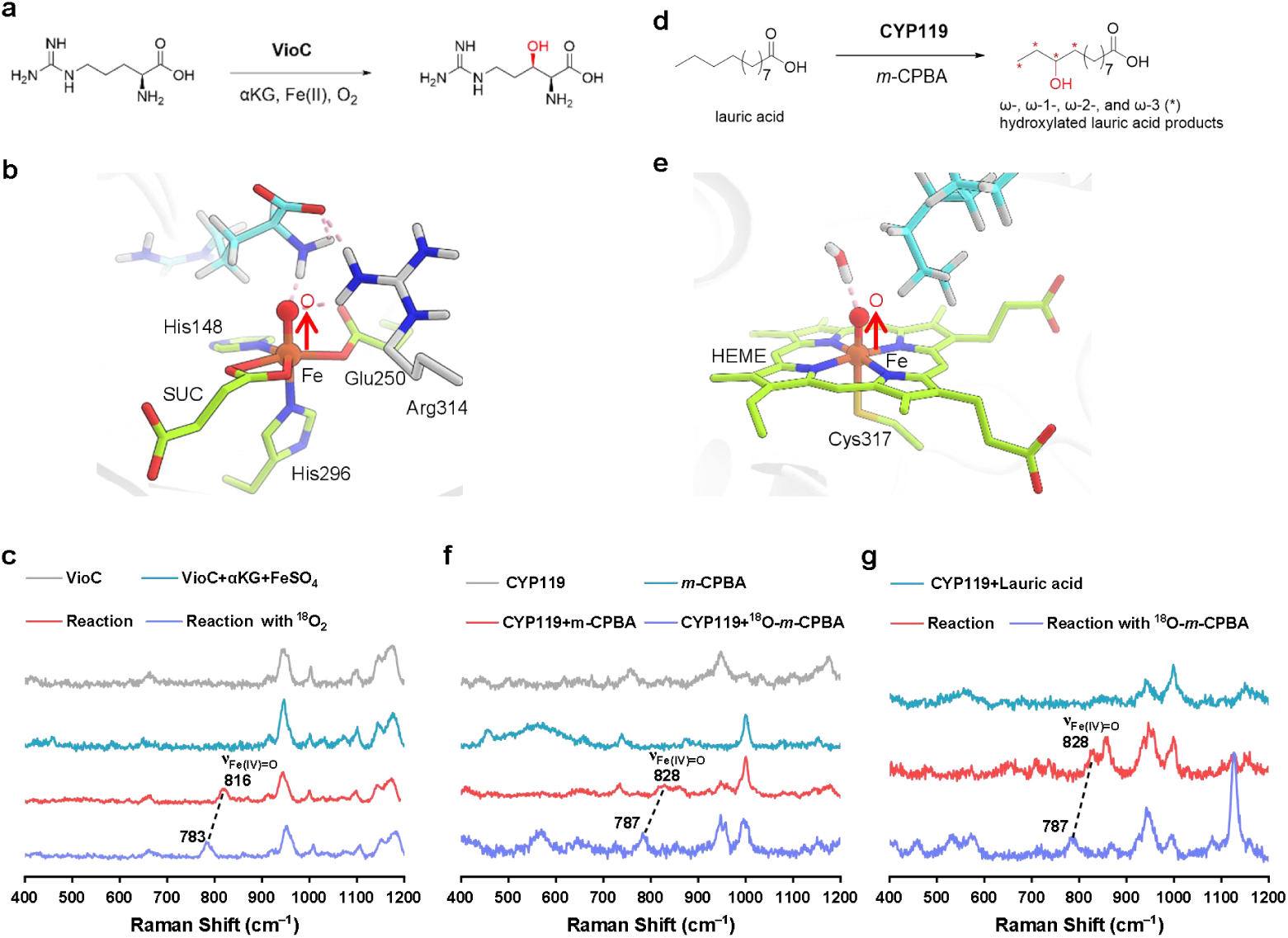
In situ SHINERS characterization of reactive Fe–O states in heme and non-heme iron enzymes. **a**, Hydroxylation of L-Arg catalyzed by VioC. **b**, QM/MM-optimized structure of the Fe(IV)=O intermediate in VioC. **c**, In situ SHINERS spectra of VioC. Reaction composition: 200 μM VioC, 1 mM L-Arg, 2 mM αKG, 200 μM FeSO4 in 150 mM Tris-HCl (pH 8.0) with 300 mM NaCl at 4 °C. For the oxygen-isotope measurement, the reaction was performed under an ^18^O_2_ atmosphere under otherwise matched conditions. **d**, Catalytic hydroxylation of lauric acid by CYP119 with *m*-CPBA. **e**, QM/MM-optimized model of CYP119 Compound I used for vibrational analysis. **f**, In situ SHINERS spectra of the CYP119/*m*-CPBA system in the absence of substrate. Reaction conditions: 20 μM CYP119, 40 μM *m*-CPBA in 100 mM potassium phosphate buffer (pH 7.0) at 4 °C. **g**, In situ SHINERS spectra of the CYP119/*m*-CPBA system in the presence of lauric acid. Reaction conditions: 20 μM CYP119, 40 μM *m*-CPBA, 40 μM lauric acid in 100 mM potassium phosphate buffer (pH 7.0) at 4 °C. The corresponding isotope spectrum was obtained by replacing unlabeled *m*-CPBA with ^18^O-labeled *m*-CPBA under otherwise identical conditions.

VioC retained catalytic competence after SHIN exposure (Supplementary Figs. 12–14). Spectra of the individual reaction components and the pre-reaction mixture were collected before initiation of catalysis. Following addition of L-arginine, a reaction-associated band appeared near 816 cm^−1^ and shifted to approximately 783 cm^−1^ under an ^18^O_2_ atmosphere (Fig. 2c). The resulting ^18^O-dependent downshift of approximately 33 cm^−1^ is close to the approximately 36 cm^−1^ shift expected from the reduced-mass isotope relation for an Fe–O vibration (Supplementary Table 1; Methods). Combined with its reaction dependence and oxygen-isotope response, these observations support assignment of the approximately 816 cm^−1^ feature to the Fe(IV)=O intermediate proposed for VioC-catalyzed hydroxylation.

For CYP119, the integrity of the resting ferric enzyme was first verified by UV–visible spectroscopy (Supplementary Fig. 16), and early-time stopped-flow measurements were used to characterize the electronic spectral evolution of the CYP119/*m*-CPBA reaction, including matched measurements in the presence of SHINs (Supplementary Figs. 17 and 18). CYP119 also retained hydroxylation activity in the presence of SHINs (Supplementary Fig. 19), demonstrating the nanoparticle environment does not abolish the reactive intermediate of CYP119. Under the SHINERS conditions, addition of *m*-CPBA produced a reaction-associated feature near 828 cm^−1^ (Fig. 2f), which shifted to approximately 787 cm^−1^ upon replacement of *m*-CPBA with ^18^O-labeled *m*-CPBA. The resulting isotope-dependent downshift of approximately 41 cm^−1^ is close to the approximately 36 cm^−1^ shift expected from the reduced-mass relation (Supplementary Table 1; Methods), supporting the Fe–O vibrational character of the ∼828 cm^−1^ band. A comparable isotope-sensitive response was observed in the presence of lauric acid (Fig. 2g), showing that the ∼828 cm^−1^ Fe–O feature is retained under substrate-turnover conditions.

To further establish the origin and assignment of the ∼828 cm^−1^ band, we combined sequential SHINERS measurements, prolonged *m*-CPBA exposure controls and QM/MM normal-mode calculations as complementary lines of evidence. Sequential SHINERS acquisition revealed a persistent ∼828 cm^−1^ response over the measurement window (Supplementary Fig. 20), consistent with time-integrated sampling of the CYP119/*m*-CPBA reaction. Prolonged exposure of CYP119 to *m*-CPBA did not reproduce the characteristic ∼828 cm^−1^ feature (Supplementary Fig. 21), arguing against its assignment to an oxidant-damaged heme species. In parallel, harmonic QM/MM calculations of the CYP119 Compound I model identified an Fe–O stretching mode in the same spectral region (Supplementary Fig. 22). Combined with the oxygen-isotope-dependent shift and the established CYP119/*m*-CPBA Compound I chemistry, these results support assignment of the ∼828 cm^−1^ band to the Fe(IV)=O vibration of CYP119 Compound I.

The VioC and CYP119 experiments extend the TauD benchmark to both non-heme and heme iron chemistry. The reproducible isotope-sensitive Fe–O vibrations observed across these systems demonstrate that in situ SHINERS can access reactive metal–oxygen intermediates across chemically distinct metalloenzyme reactions, including the Fe(IV)=O unit of P450 Compound I.

### Evidence for an (R)HN–Fe(IV)=O intermediate supports dynamic coordination in TqaL_Nc_

Beyond classical hydroxylation chemistry, Fe(II)/αKG-dependent enzymes catalyze a diverse range of noncanonical oxidative transformations^10^. A central unresolved question in this field concerns how highly reactive ferryl intermediates are selectively redirected away from hydroxylation toward alternative bond-forming reactions. TqaL_Nc_ catalyzes an unusual aziridine-forming C–N coupling reaction from L-valine (L-Val) (Fig. 3a)^55,56^. Our recent computational work proposed that TqaL_Nc_ utilizes a dynamic coordination mechanism, involving both a conformational switch of the Fe(IV)=O intermediate and dynamic coordination of the substrate amino group to the iron center^57^. This process leads to the formation of a key intermediate, (R)HN–Fe(IV)=O, which promotes the following C–H activation and C–N coupling reactions (Fig. 3b). However, direct experimental evidence for such a species had remained unavailable.

**Fig. 3.**
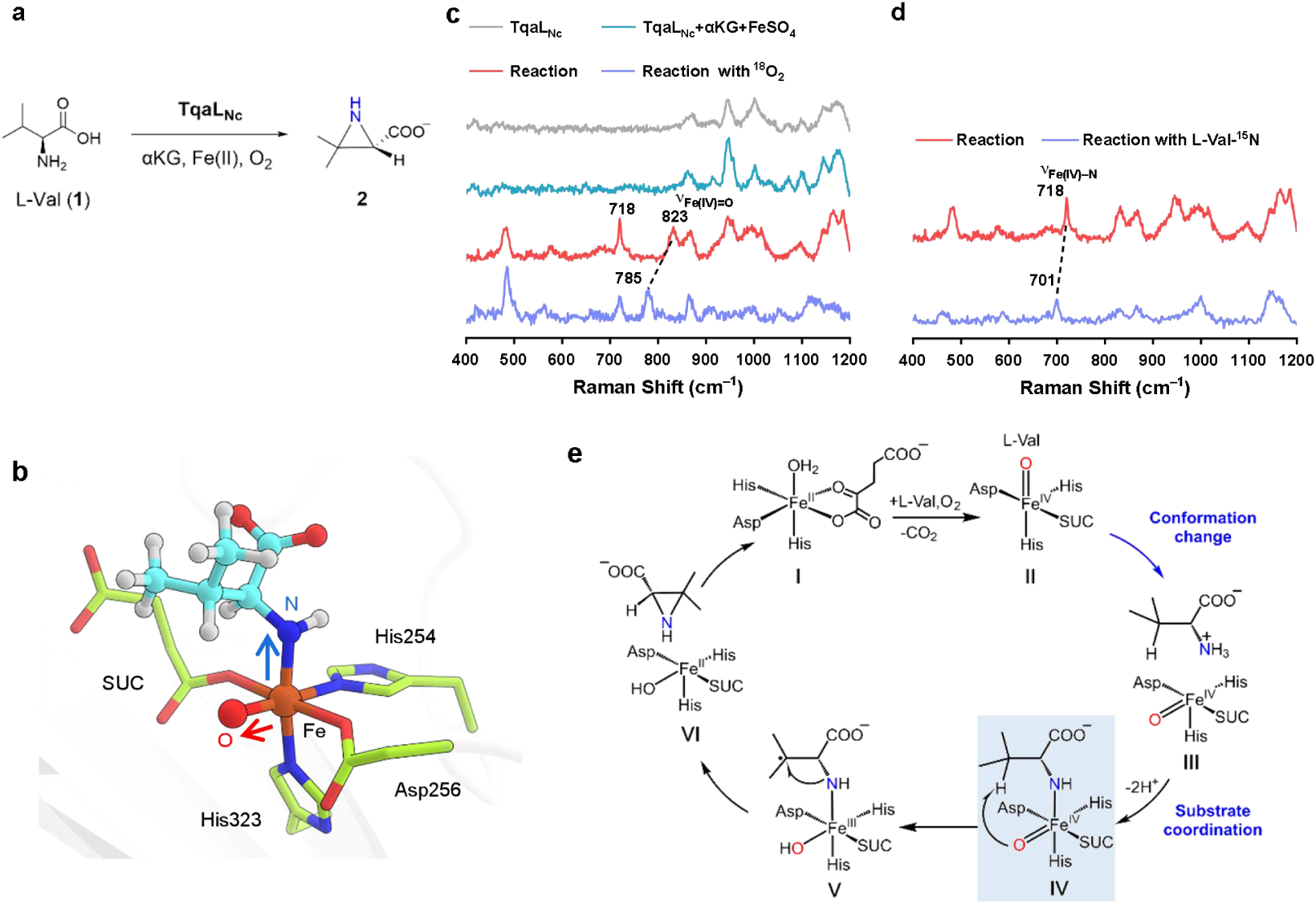
In situ SHINERS evidence supporting an (R)HN–Fe(IV)=O coordination state in TqaL_Nc_. **a**, The C–N coupling reaction catalyzed by TqaL_Nc_. **b**, The QM/MM-optimized structure of the proposed (R)HN–Fe(IV)=O intermediate. **c**, In situ SHINERS spectra of TqaL_Nc_. Reaction composition: 200 μM TqaL_Nc_, 1 mM L-Val, 2 mM αKG, 200 μM FeSO_4_, 150 mM Tris-HCl (pH 8.0) with 300 mM NaCl at 4 °C. For the oxygen-isotope measurement, the reaction was performed under an ^18^O_2_ atmosphere under otherwise matched conditions. **d**, In situ SHINERS spectra of TqaL_Nc_ reaction with L-Val-^15^N. For the nitrogen-isotope measurement, unlabeled L-Val was replaced with ^15^N-labeled L-Val at the same final concentration under otherwise identical reaction conditions. **e,** Proposed catalytic cycle for TqaL_Nc_, illustrating the dynamic coordination mechanism that enables aziridine formation. SUC: succinate.

TqaL_Nc_ retained catalytic activity in the presence of SHINs (Supplementary Figs. 24–26), demonstrating compatibility of the SHINERS configuration with the aziridine-forming reaction. In the complete reaction, two reaction-associated features were observed near 718 and 823 cm^−1^ (Fig. 3c). Under an ^18^O_2_ atmosphere, the approximately 823 cm^−1^ band shifted to approximately 785 cm^−1^, corresponding to a downshift of approximately 38 cm^−1^ and supporting assignment to an Fe(IV)=O vibration (Fig. 3c). In contrast, the approximately 718 cm^−1^ feature was insensitive to ^18^O substitution but shifted to approximately 701 cm^−1^ when L-Val-^15^N was used (Fig. 3d), corresponding to a ^15^N-dependent downshift of approximately 17 cm^−1^. The distinct oxygen- and nitrogen-isotope responses identify the two bands as Fe–O- and substrate-N-associated vibrational modes, respectively.

Harmonic QM/MM normal-mode calculations for the proposed (R)HN–Fe(IV)=O structure yielded modes containing Fe–N and Fe–O stretching character near 693 and 827 cm^−1^, respectively (Supplementary Fig. 29). The experimentally observed ^18^O- and ^15^N-dependent shifts are directionally consistent with simple reduced-mass estimates (Supplementary Table 1; Methods). The isotope responses and QM/MM-calculated mode character therefore provide convergent vibrational evidence for the proposed (R)HN–Fe(IV)=O coordination state.

In the previously proposed mechanism of TqaL_Nc_ (Fig. 3e)^57^, αKG-assisted oxygen activation first generates a ferryl Fe(IV)=O species (I→II), followed by reorientation of the oxo unit from an axial to an equatorial configuration (II→III). Subsequent deprotonation and coordination of the substrate amino group to the iron center yields the NH-ligated ferryl state, (R)HN–Fe(IV)=O (III→IV). This coordination state is proposed to facilitate C–H activation (IV→V), generating a substrate radical that subsequently undergoes intramolecular C–N bond formation (V→VI).

The ^18^O- and ^15^N-sensitive shifts and QM/MM analysis support formation of the proposed (R)HN–Fe(IV)=O state and substrate-nitrogen coordination to the ferryl center. This experimental evidence anchors the dynamic-coordination model and explains how TqaL_Nc_ redirects high-valent iron reactivity from conventional hydroxylation toward aziridine-forming C–N coupling. More broadly, the results show that SHINERS can resolve not only metal–oxygen intermediates but also mechanistically important changes in metal–substrate coordination.

### Interfacial basis of SHINERS sensitivity in metalloenzyme systems

To understand how the SHIN interface facilitates vibrational measurements of metalloenzymes, we examined enzyme–SHIN interactions by scanning electron microscopy (SEM) and molecular dynamics (MD) simulations. SEM images showed association of TauD with the silica-coated nanoparticle surface (Fig. 4a). MD simulations further identified stable adsorption configurations in which polar and charged surface residues form hydrogen-bonding and salt-bridge interactions with amorphous silica (Fig. 4b and Supplementary Figs. 31–34). Similar adsorption configurations were obtained for the other metalloenzymes examined, indicating that the silica shell provides a general interfacial environment that positions enzyme molecules close to the plasmonic surface.

**Fig. 4.**
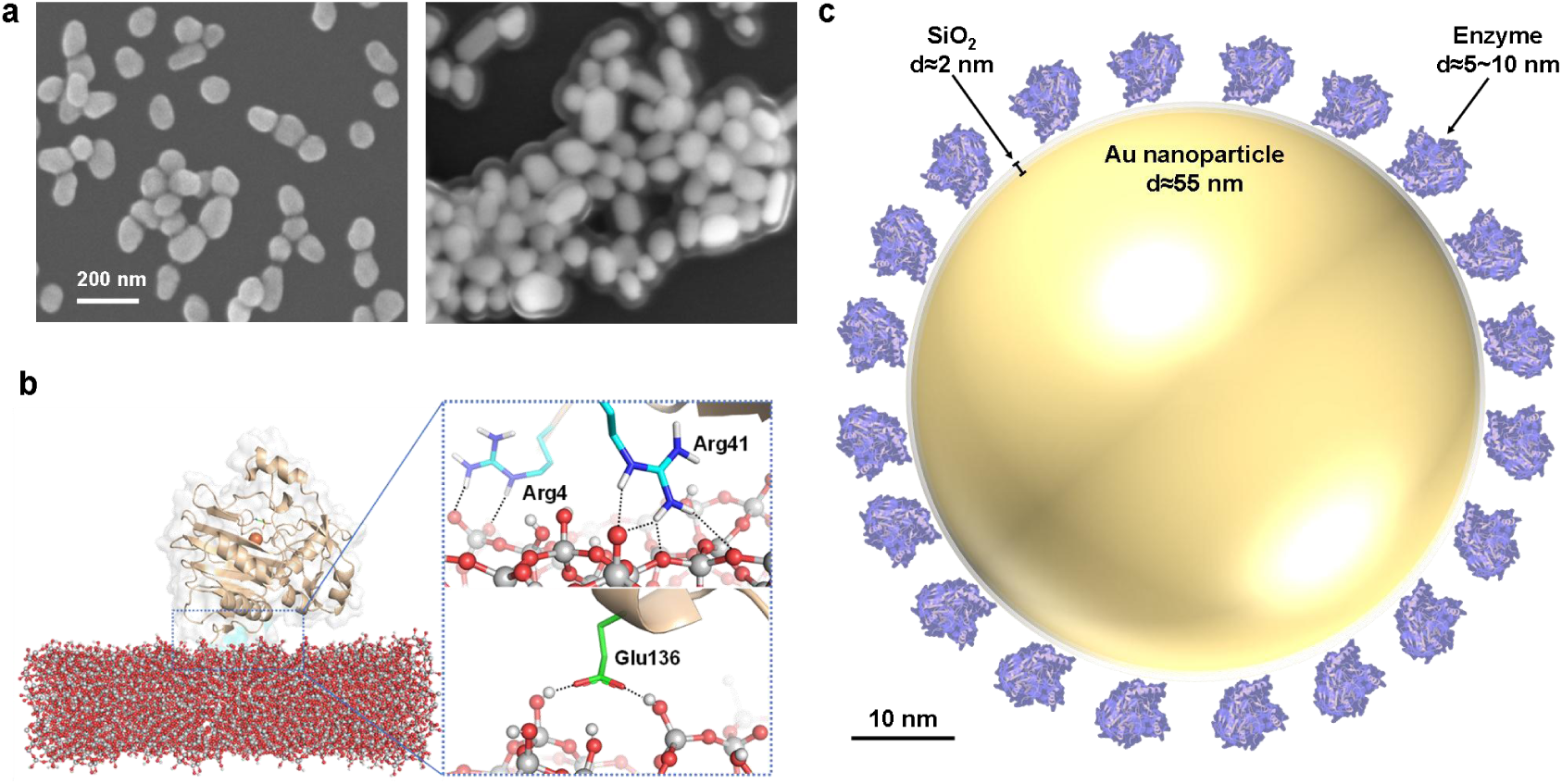
Interfacial association of metalloenzymes with SHINs. **a**, Scanning electron microscopy (SEM) images of SHINs before (left) and after incubation with TauD (right), showing protein association with the silica-coated nanoparticle surface. **b**, Representative MD-derived adsorption configuration of TauD on an amorphous silica surface, highlighting hydrogen-bonding and salt-bridge interactions between protein surface residues and silica. **c**, Schematic representation of enzyme association with Au@SiO_2_ SHINs and the relative dimensions of the nanoparticles and adsorbed proteins.

Such enzyme–silica association places portions of the protein within the electromagnetic near-field generated by the Au core while maintaining physical separation from direct contact with the metal surface (Fig. 4c). This interfacial proximity provides a physical basis for enhanced Raman sampling of weak protein and metal–ligand vibrational modes, with the observed signal determined by the local enzyme–SHIN configuration.

The combined SEM and MD results establish an interfacial model for SHIN-assisted Raman interrogation of metalloenzymes under aqueous reaction conditions. By bringing enzyme molecules into the plasmonically enhanced near-field while preserving a chemically isolating silica interface, the SHIN architecture provides access to weak metal–oxygen and metal–substrate vibrational signatures during enzymatic reactions.

## Discussion

This study establishes in situ SHINERS as a powerful vibrational approach for visualizing reactive metal–ligand states associated with metalloenzyme intermediates under aqueous reaction conditions. Across the heme and non-heme iron enzymes examined here, SHINERS resolves isotope-sensitive metal–oxygen and metal–substrate vibrations. In TauD and VioC, the measurements provide vibrational characterization of ferryl intermediates, while the CYP119 results identify the Fe(IV)=O vibration of P450 Compound I generated during *m*-CPBA-driven chemistry. These results extend SHINERS from established applications in interfacial catalysis to reactive coordination states in enzymatic systems.

The capability of SHINERS arises from its interfacial architecture. Association of metalloenzymes with the silica-coated nanoparticles places protein molecules within the plasmonically enhanced near-field while maintaining physical separation from direct contact with the Au core. This configuration facilitates Raman sampling of weak metal–ligand modes without requiring electronic resonance of the probed vibration. SHINERS adds direct vibrational information to the kinetic, electronic and structural observables provided by established spectroscopic and structural methods.

SHINERS also provides vibrational evidence supporting the dynamic-coordination mechanism underlying selective C–N bond formation in TqaL_Nc_. The complementary ^18^O- and ^15^N-dependent shifts, together with QM/MM normal-mode analysis, support formation of the proposed (R)HN– Fe(IV)=O state and directly connect substrate-nitrogen coordination to the ferryl center. In this framework, substrate-nitrogen coordination alters the primary coordination environment of Fe(IV)=O, redirecting its reactivity from conventional hydroxylation toward selective aziridine-forming C–N coupling^58,59^. Thus, the TqaL_Nc_ results show how SHINERS can move beyond canonical metal–oxygen intermediates to reveal coordination changes underlying noncanonical enzymatic reactivity.

More broadly, these findings expand experimental access to the metal–ligand vibrational signatures of reactive intermediates in metalloenzyme catalysis. Interrogating both metal–oxygen and metal–substrate coordination under aqueous reaction conditions directly links transient coordination chemistry to enzymatic reactivity and selectivity. Future integration of SHINERS with rapid-mixing or continuous-flow reaction initiation, electrochemical triggering, and coordinated electronic and vibrational measurements could add a defined temporal dimension to this approach and enable direct correlation between vibrational structure and transient-state kinetics.

## Materials and methods

Unless otherwise noted, chemicals and reagents were purchased from Sigma-Aldrich, Sangon Biotech, Energy Chemical, Acmec or Solarbio. LB medium, 20× PBS Buffer, TureColor Marker and Modified Bradford Protein Assay Kit were obtained from Sangon Biotech. Kanamycin and isopropyl β-D-1-thiogalactopyranoside (IPTG) were supplied by Solarbio. Lauric acid, αKG, taurine, Tris, Li_2_CO_3_ and dansyl chloride were procured from Acmec. L-Arginine, L-Valine and δ-aminolevulinic acid were provided by Energy Chemical. Trifluoroacetic acid and acetonitrile were acquired from Sigma-Aldrich. L-Valine-^15^N was supplied by Medlife, L-Valine-^2^d8 by IsoReag, and ^18^O_2_ gas (≥99% isotopic purity) by Wuhan Isotope Technology. ^18^O-labeled *m*-CPBA was prepared from 3-chlorobenzoic acid and H_2_-^18^O_2_ using a reported procedure^60^.

### Synthesis of the SHINs and structural characterization

Prior to the fabrication of Au@SiO_2_ shell-isolated nanoparticles (SHINs) for Shell-Isolated Nanoparticles enhanced Raman Spectroscopy (SHINERS), 55 nm Au nanoparticles (AuNPs) were synthesized according to the Frens method^61^. Specifically, 200 mL of 0.01% chloroauric acid (HAuCl_4_) aqueous solution was boiled and stirred in a round-bottom flask, followed by the addition of 1.4 mL of 1% sodium citrate solution. The reaction was maintained for approximately 30 minutes. After the solution cooled to room temperature, 55 nm Au nanoparticles were obtained.

The subsequent preparation of Au@SiO_2_ shell-isolated nanoparticles (SHINs) involved the following steps: Initially, 0.4 mL of 1 mM 3-aminopropyltrimethoxysilane (APTMS) solution was added dropwise to 30 mL of the freshly prepared 55 nm Au nanoparticle solution, and the mixture was stirred for 15 min. Subsequently, 3.2 mL of sodium silicate (Na_2_SiO_3_) solution, which had been hydrolyzed for 15 min, was poured into the mixture, and stirring was continued for an additional 5 min. The mixture was then heated and stirred in a water bath at 98 ℃ for 20 min. After cooling the solution in an ice bath, the mixture was centrifuged three times at 5500 rpm to obtain Au@SiO_2_ shell-isolated nanoparticles. The concentrated shell-isolated nanoparticles were further diluted to 0.4 mL of water for subsequent Raman testing.

The plasmonic properties of Au@SiO_2_ SHINs with Au cores and ultrathin silica shells have been extensively characterized in previous SHINERS studies^40,42,46^. For Au cores of approximately 50– 60 nm, comparable to the approximately 55 nm Au cores used here, the characteristic localized surface plasmon response is typically observed in the visible region at approximately 530–550 nm. Formation of an ultrathin silica shell largely preserves the plasmonic response of the Au core while allowing the enhanced electromagnetic near-field to extend beyond the dielectric shell^40,42,46^.

The morphology of the SHINs was examined by scanning electron microscopy (SEM; Zeiss GeminiSEM 500) and high-resolution transmission electron microscopy (TEM; JEOL JEM-2100plus and JEOL JEM-F200 equipment at an accelerating voltage of 200 kV). Prior to imaging, a 3 μL aliquot of the diluted SHINs was drop-cast onto a silicon wafer and dried at ambient conditions. To prepare the SHINs-enzyme complexes, 100 μL of diluted SHINs was mixed with 100 μL of the enzyme solution (the “Preparation of Metalloenzymes” section) to yield a uniform turbid suspension. A 3 μL aliquot of this resultant mixture was then deposited onto a silicon wafer and air-dried.

### Preparation of metalloenzymes

The genes encoding CYP119, TauD, VioC, or TqaL_Nc_ (amino acid sequences are provided in the Supplementary Information) were codon-optimized for *E. coli* expression and synthesized by GenScript (Nanjing, China). Each gene was subsequently cloned between the *Nde* I and *Xho* I restriction sites of the pET-28a(+) vector to generate an N-terminal His_6_-tagged construct. The resulting recombinant plasmids were transformed into *E. coli* BL21(DE3) competent cells for protein production^62^. Transformants were grown at 37 °C in LB medium supplemented with 40 mg/L kanamycin. Protein expression for all targets was induced at an OD_600_ of 0.6–0.8, with induction performed as follows: CYP119 was induced with 0.5 mM IPTG and 0.3 mM δ-aminolevulinic acid (5-ALA), followed by 16 h of incubation at 30 °C^63^; TauD was induced with 0.5 mM IPTG for 16 h at 25 °C^64^; for both VioC and TqaL_Nc_, cultures were cooled on ice for 30 min prior to induction with 0.2 mM IPTG and subsequent incubation for 18 h at 16 °C^56,65^.

Following induction, cells were harvested by centrifugation and resuspended in lysis buffer (20 mM Na_3_PO_4_, 500 mM NaCl, 20 mM imidazole, pH 7.4). The suspensions were sonicated on ice. For CYP119, the sonicated lysate was incubated at 65 °C for 1 h before centrifugation, using the thermostability of CYP119 to precipitate heat-labile host proteins^63^. The heat-treated CYP119 lysate and the untreated lysates of the other enzymes were centrifuged at 10,000 rpm for 15 min to remove insoluble material. Each supernatant was loaded onto Ni-NTA agarose, washed with lysis buffer containing 20 mM imidazole, and eluted with buffer containing 300 mM imidazole. The eluate was then desalted using a PD-10 desalting column (Cytiva)^66^. Protein purity was assessed by SDS-PAGE. Concentrations of TauD, VioC, and TqaLNc were determined by a modified Bradford assay. CYP119 concentration was determined from the reduced pyridine hemochromogen spectrum of heme b^67^.

### Metalloenzyme activity assays

CYP119 assays were performed in 50 μL reactions containing 100 mM potassium phosphate (pH 7.0), 1 mM *m*-CPBA, 1 mM lauric acid, and 20 μM ferric CYP119 ^14,68^. Reactions were incubated at 4 °C for 1 h and stopped by adding 5 μL of 1 M HCl. Each acidified sample was extracted three times with 200 μL of ethyl acetate; the organic layers were combined and evaporated to dryness.

For HPLC analysis of CYP119 products, the dried ethyl-acetate extract was derivatized to form p-bromophenacyl (PBAB; also abbreviated p-BPBr) esters using p-bromophenacyl bromide and 18-crown-6, adapted from established fatty-acid and hydroxy-fatty-acid derivatization procedures^69,70^. The residue was dissolved in 67.6 μL of acetonitrile, followed by 4.4 μL of 25 mM KOH and mixing for 2 min. Then, 10 μL of 1 mM 18-crown-6 was added and the sample was mixed for 3 min. A 15 μL aliquot of 10 mM p-BPBr and a further 103 μL of acetonitrile were added. The sealed sample was agitated at 80 °C for 15 min, cooled to room temperature, and centrifuged; the clear supernatant was analyzed by HPLC. A Waters HPLC system equipped with a ZORBAX Eclipse Plus C18 column (250 × 4.6 mm, 5 μm; Agilent) was used with 0.1% (v/v) trifluoroacetic acid in water (A) and acetonitrile (B). The gradient was 70-90% B over 0-15 min, 90% B over 15-18 min, and return to 70% B over 18-23 min at 1 mL/min. The p-bromophenacyl derivatives were detected at 254 nm.

Assays for Fe(II)/αKG-dependent enzymes (TauD, VioC, and TqaL_Nc_) were performed as described previously^56^. The standard reaction system contained 150 mM Tris-HCl (pH 8.0), 300 mM NaCl, 0.2 mM FeSO_4_, 2 mM αKG, 1 mM substrate, and 20 μM enzyme. Reactions with TauD and VioC were conducted at 30 °C for 1 h, whereas TqaL_Nc_ was assayed at 20 °C. To investigate the effect of nanoparticles, each enzyme was pre-incubated with AuNPs or SHINs for 10 min before the reaction was initiated under its respective standard assay temperature (30 °C for TauD and VioC and 20 °C for TqaL_Nc_). The impact of low temperature on catalytic activity was assessed by performing reactions at 4 °C for 1 h. For the kinetic isotope effect (KIE) studies of TqaL_Nc_, reactions contained 40 μM TqaL_Nc_, 2 mM αKG, 200 μM FeSO_4_, 1 mM L-Val or L-Val-^2^d8, 150 mM Tris-HCl (pH 8.0), and 300 mM NaCl and were performed at 20 °C.

Prior to HPLC analysis, the reaction products of TauD, VioC, and TqaL_Nc_ were derivatized with dansyl chloride^55^. Specifically, a 50 μL aliquot of the reaction mixture was mixed with 50 μL of 80 mM Li_2_CO_3_, 45 μL of acetonitrile, 25 μL of water, and 30 μL of 10 mM dansyl chloride (in acetone). The mixture was then incubated at room temperature for 1 h. The derivatized samples were analyzed using the same HPLC system and column as described above, with the following gradient program: 0–15 min, 10% to 90% B; 15–18 min, 90% to 10% B; 18–23 min, 10% B. Detection was performed at 220 nm.

For LC-MS analysis, samples were injected into an Agilent 1290 system coupled to a 6546 Q-TOF mass spectrometer and equipped with a ZORBAX Eclipse Plus C18 column (250 mm × 4.6 mm, 5 μm; Agilent). Separation was achieved using the same gradient program as described above at a flow rate of 1 mL/min.

### In situ SHINERS measurements

In the present study, “in situ” refers to acquisition of Raman spectra from the reacting aqueous enzyme mixture while it remains in contact with the SHIN-coated solid/liquid interface. Unless otherwise specified, spectra were collected with a 120 s integration time. The confocal excitation and collection volume was focused on the SHIN-rich interfacial region, where diffusion, adsorption and interfacial mass transport can contribute to the measured temporal response^43–47^.

Raman spectra were acquired on a HORIBA XploRA confocal micro-Raman system equipped with a 785 nm semiconductor laser operated at approximately 6 mW^47^. A 50× long-working-distance objective with an 8 mm working distance was used, and the spectrometer was calibrated using a silicon wafer and perchlorate solution. For sample preparation, a 3 μL aliquot of diluted Au@SiO_2_ SHINs was drop-cast onto a glassy-carbon electrode and air-dried at room temperature. The enzyme-containing aqueous reaction mixture was subsequently introduced into the sealed Raman cell and brought into contact with the SHIN-coated surface. The confocal excitation and collection volume was focused on the SHIN-rich solid/liquid interfacial region (Supplementary Fig. 3). Diffusion, adsorption, and interfacial mass transport may contribute to the measured temporal response.

The measurement workflow comprised catalytic-competence controls, spectra of individual reaction components, acquisition of a pre-reaction spectrum, collection after reaction initiation, isotope-substitution measurements, and comparison with QM/MM normal-mode calculations.

For CYP119 SHINERS measurements, reference spectra of CYP119, *m*-CPBA, lauric acid, and the relevant pre-reaction mixtures were recorded before reaction initiation. All CYP119 SHINERS measurements were performed in 100 mM potassium phosphate buffer (pH 7.0) at 4 °C. Unless otherwise specified, reactions were initiated by addition of *m*-CPBA, and Raman acquisition was started immediately after reaction initiation. For the no-substrate reaction, the final concentrations were 20 μM CYP119 and 40 μM *m*-CPBA. For turnover measurements, the reaction contained 20 μM CYP119, 40 μM lauric acid, and 40 μM *m*-CPBA. Oxygen-isotope measurements were performed by replacing unlabeled *m*-CPBA with ^18^O-labeled *m*-CPBA under otherwise identical conditions. For prolonged-oxidant controls, 20 μM CYP119 was mixed with 40 μM *m*-CPBA and incubated for 5 or 10 min at 4 °C before SHINERS measurement.

A separate sequential-acquisition experiment was performed to examine the relationship between reaction initiation and the time-integrated SHINERS signal. In this experiment only, Raman acquisition was initiated before *m*-CPBA addition, and the oxidant was injected directly into the Raman cell during ongoing spectral integration. Consecutive spectra were then collected over 0– 120, 120–240, and 240–360 s after reaction initiation.

For SHINERS measurements of TauD, VioC, and TqaL_Nc_, spectra of the individual reaction components and the corresponding pre-reaction mixtures were recorded first. Reactions were then initiated by addition of the corresponding substrate, and Raman acquisition was started immediately after reaction initiation. The pre-reaction mixture contained 200 μM enzyme, 200 μM FeSO₄, and 2 mM αKG in 150 mM Tris-HCl (pH 8.0) containing 300 mM NaCl. Reactions were initiated by addition of the corresponding substrate to a final concentration of 1 mM (taurine for TauD, L-Arg for VioC, and L-Val for TqaL_Nc_). SHINERS measurements were performed at 4 °C. Oxygen-isotope experiments were performed under an ^18^O_2_ atmosphere under otherwise matched conditions. For TqaL_Nc_, nitrogen-isotope measurements were performed by replacing unlabeled L-Val with ^15^N-labeled L-Val (^15^N-L-Val) at the same final substrate concentration.

### Spectral processing

Raw Raman spectra were retained for all measurements. Within each matched comparison, the same baseline-correction procedure and parameter settings were applied over the same spectral range. Baseline correction was used only to remove slowly varying background contributions while minimizing distortion of genuine Raman bands. The fitted baseline was constrained to follow the broad spectral background rather than individual Raman peaks or broad vibrational features.

When smoothing was applied, identical smoothing parameters were used for all spectra within the corresponding comparison, and matched isotope spectra were treated using the same normalization procedure. No local anchor points or manual baseline adjustments were introduced within the assigned Fe–O or Fe–N spectral regions. Raw and processed spectra were evaluated in parallel, and spectral assignments were based primarily on reaction dependence, reproducible isotope-dependent peak positions, matched controls, and independent replicate measurements rather than on differences in processed peak intensity. Representative unprocessed spectra are provided for the TauD reaction in Supplementary Fig. 10 and for the TqaL_Nc_ reaction in Supplementary Figs. 27 and 28.

### Resonance Raman spectroscopy and SERS measurements

Resonance Raman and surface-enhanced Raman spectroscopy (SERS) analyses were performed for the Fe(II)/αKG-dependent enzyme TauD. The SHINERS workflow was used with the following modifications. For resonance Raman measurements, the key modifications were the concentration of the reaction system (400 μM FeSO_4_ and 400 μM TauD), the use of a 325 nm laser excitation source and a measurement temperature of −40 °C. All other experimental parameters remained unchanged. For SERS sample preparation, a 3 μL aliquot of the diluted SHINs solution was replaced with silica-uncoated Au nanoparticles (AuNPs). This AuNP solution was drop-cast onto a glassy carbon electrode and allowed to air-dry at room temperature. All other preparation and measurement conditions were kept consistent.

For the TauD resonance-Raman and uncoated-Au controls, spectra were collected for TauD alone, for the pre-reaction mixture containing TauD, FeSO_4_, and alpha-ketoglutarate, and after taurine addition. Paired isotope spectra were acquired and processed using matched settings. In the experiments, the shaded region denotes the expected ferryl spectral region and does not indicate an independently resolved Fe(IV)-oxo band.

### Stopped-flow absorption spectroscopy measurements

Stopped-flow absorption measurements were performed using an Applied Photophysics SX20 spectrophotometer at 4 °C. Equal volumes of CYP119 and *m*-CPBA solutions were rapidly mixed to give final concentrations of 10 μM CYP119 and 20 μM *m*-CPBA in 100 mM potassium phosphate buffer (pH 7.0). Photodiode-array spectra were collected during the earliest accessible reaction period, and representative kinetic traces were monitored at approximately 366, 416, and 690 nm. Matched measurements were performed in the presence of SHINs to characterize the early-time electronic spectral evolution of the CYP119/*m*-CPBA system and evaluate the influence of SHINs on this behavior.

### System setup and MD simulations

#### (1) Classical MD Simulations of Proteins in Aqueous Solution

To obtain reliable structural models for Raman spectra calculations, we performed classical molecular dynamics (MD) simulations for each enzyme in their respective intermediate states. The crystal structures of CYP119 (PDB: 1IO7), TauD (PDB: 1GQW) and VioC (PDB:6ALR) were used to build the model systems. Substrates were docked into the active site separately using the AutoDock Vina tool. We assigned the protonation states of titratable residues (His, Glu, Asp) on the basis of p*K*_a_ values using PROPKA and a visual inspection of local H-bonding networks. All Glu and Asp residues were deprotonated. The missing hydrogen atoms were added by the LEAP module in Amber 18. The general AMBER force field (GAFF) was used for the substrates. The partial atomic charges were fitted using the RESP method, using the HF/6-31G* level of theory. The force fields for Compound I (Cpd I) and the Fe(IV)=O species were parametrized using MCPB^71^. Several Na^+^ ions were added to the surface of the protein to neutralize the total charge of the systems. Finally, the neutral systems were solvated in a cube of TIP3P waters within a 15 Å water layer. The Amber ff14SB force field was used for the protein residues.

After setup, the systems were minimized by a combined steepest descent and conjugate gradient method. Then the systems were heated from 0 to 300 K under a canonical ensemble for 0.2 ns with a weak restraint of 15 kcal/(mol·Å^2^). To achieve a uniform density after heating dynamics, 1 ns of density equilibration was performed under the NPT ensemble at the target temperature of 300 K and target pressure of 1.0 atm. Afterward, we removed all restraints on the protein and further equilibrated the system for 4 ns under the NPT ensemble to equilibrate pressure and temperature using Langevin thermostat and Berendsen barostat. Finally, a production MD run was conducted for 200 ns. The covalent bonds containing hydrogen were constrained using SHAKE. Representative snapshots from the MD trajectory were used as starting structures for the QM/MM calculations. For TqaL_Nc_, the MD simulations have been conducted in our previous work^57^.

#### (2) MD Simulations of Protein Adsorption on Amorphous Silica

To investigate the adsorption of four proteins (TauD, VioC, CYP119, and TqaL_Nc_) onto amorphous silica, we performed classical molecular dynamics (MD) simulations for each enzyme in an aqueous solution. The INTERFACE force field^72,73^ was employed to describe the SiO_2_ slab. The amorphous silica slab model was constructed using the structural coordinates reported by Emami et al.^74^, and the SiO_2_ surface was expanded to a size of 80 Å × 80 Å to avoid periodic boundary effects. Considering the hydration behavior of SiO_2_ in a neutral aqueous environment, 18% of the surface oxygen atoms were left in a non-hydrated state. After the SiO_2_ surface was constructed, each protein was positioned directly above it using Packmol^75^, with a geometric center separation of 50 Å between the protein and the silica slab. Protein force field parameters were assigned as detailed in Section 5.1. To systematically sample initial adsorption orientations, multiple rotated configurations were generated for each protein: first, by rotating the protein around the x-axis by 90°, 180°, and 270° while constraining the y- and z-axes, resulting in four configurations designated Rx (0°) Ry(0°)-box1, Rx (90°) Ry(0°)-box2, Rx(180°) Ry(0°)-box3, and Rx(270°) Ry(0°)-box4. Subsequently, starting from the Rx(0°) Ry(0°)-box1 orientation, rotations around the y-axis by 90°, 180°, and 270° were performed with the x- and z-axes fixed, yielding three additional structures: Rx(0°) Ry(90°)-box5, Rx(0°) Ry(180°)-box6, and Rx(0°) Ry(270°)-box7. In total, these seven distinct initial configurations per protein were used in subsequent molecular dynamics simulations to study protein adsorption onto amorphous silica in an aqueous environment.

Following the initial setup, each system was solvated in a rectangular water box measuring 100 Å × 100 Å × 135 Å, filled with TIP3P water molecules. Na⁺ ions were then added to neutralize the total charge of the simulation system. During all simulations, the silica slab was held fixed by applying harmonic positional restraints with a force constant of 50 kcal/(mol·Å^2^) to all silicon atoms. Finally, a 200 ns NPT production simulation was performed for each box, from which the corresponding stable adsorbed states were extracted for analysis.

### Raman spectra calculations

Representative snapshots from the production MD trajectory in Section 5.1 were selected for Raman spectra calculations using a QM/MM approach. All QM/MM calculations were performed using ChemShell^76^, combining ORCA as the QM code and DL_POLY^77^ as the MM code. Residues within 12.0 Å of the Fe atom were included in the active region. For TauD, the QM region contains the Fe(IV)=O species, substrates, succinate, the truncated residues D197, H195, H246 coordinated to Fe atom, truncated residues N91, K261 and two water molecules that form H-bonding interactions with the substrate. For CYP119, the QM region contains the substrate, Cpd I and a water molecule which is coordinated to the O atom in Cpd I. For TqaL_Nc_, the QM region contains the Fe(IV)=O species, substrates, succinate, the truncated residues D256, H254, H323 coordinated to Fe atom, and three water molecules that form H-bonding interactions with the substrate and Fe(IV)=O species. The electronic embedding^78^ scheme was employed to include the polarizing effect of the enzyme environment on the QM region.

Geometries were optimized in the doublet state for CYP119 and quintet state for TauD and TqaL_Nc_ using the BP86 functional with def2-TZVP basis sets on Fe, N, and O, while def2-SVP basis sets were used for C and H. The RI-J approximation was used with def2/J basis sets. For the optimizations, TightOpt settings with tighter integration grids (Grid6) were used. Dispersion corrections were included using D3BJ. Raman spectra were calculated using the same level of theory as the geometry optimizations.

Harmonic normal-mode calculations were performed for the standard-isotope intermediate models to evaluate vibrational frequencies and mode character. No isotope-substituted QM/MM calculations were performed; the ^18^O- or ^15^N shifts in Supplementary Table 1 were instead estimated using the reduced-mass relation described below. No anharmonic correction or empirical frequency-scaling factor was applied. Accordingly, the calculations were used primarily to evaluate mode character rather than exact agreement between calculated and experimental frequencies.

### Estimation of isotope-induced vibrational frequency shifts

Oxygen-isotope measurements for TauD, VioC, and TqaL_Nc_ were performed under an ^18^O_2_ atmosphere under otherwise matched reaction conditions (Figs. 1d, 2c, and 3c). For CYP119, oxygen-isotope substitution was achieved by replacing unlabeled *m*-CPBA with ^18^O-labeled *m*-CPBA (Fig. 2f, 2g). The reproducible shifts to lower wavenumber upon ^18^O substitution were used to identify vibrational modes containing substantial oxygen-associated character. For TqaL_Nc_, a corresponding ^15^N-isotope experiment was performed by replacing L-Val with L-Val-^15^N (Fig. 3d), allowing identification of a substrate-nitrogen-associated vibrational mode.

The theoretical isotope shifts reported in Supplementary Table 1 were calculated from these ratios and the experimentally observed unlabeled frequencies. These isotope-shift estimates are independent of the QM/MM normal-mode calculations. These reduced-mass estimates are used as simplified reference values for the expected direction and approximate magnitude of isotope-dependent shifts. The experimentally observed vibrations are normal modes of a polyatomic metal– ligand environment; the reduced-mass estimates are therefore not interpreted as evidence that the corresponding bands represent pure isolated diatomic oscillators.

For intermediate species involving *O or *N, the relative Raman frequency shift (γ) observed in the ^18^O_2_ or L-Val-^15^N experiments can be approximated using the mass formula:

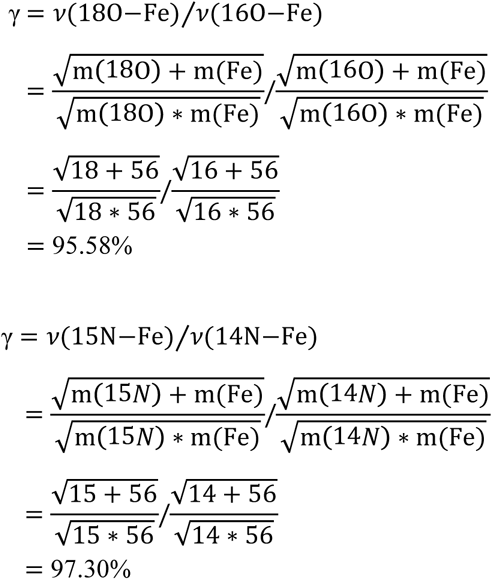

## Data availability

All data are available in the main text or Supplementary Information.

## Acknowledgments

We are grateful to Prof. Zhong-Qun Tian (Xiamen University) for his valuable suggestions and discussions. We thank Prof. Haohua Huo and Zhen Xu (Xiamen University) for their help with the synthesis of ^18^O-labeled *m*-CPBA. We thank Yuan Wang (Xiamen University) for her assistance with the LC–MS measurements. We also thank Prof. Wenjing Hong and Aihui Zhang (Xiamen University) for providing access to instrumentation. This work was supported by the Scientific Research Innovation Capability Support Project for Young Faculty (ZYGXONJSKYCXNLZCXM-B6 to B.W.), the National Natural Science Foundation of China (22525042 and T2293692 to J.-F.L., 22122305 to B.W., 22303043 to X.Z., and 22403016 to J.L.), and State Key Laboratory Program of Physical Chemistry of Solid Surfaces, Xiamen University (202430 to X.Z.).

## Author contributions

B.W. and J.-F.L. conceived the project and directed the research. L.L. performed the genetic and biochemical experiments. Q.-Q.L., Z.-H.C. and L.L. carried out the Raman experiments. X.Z. undertook the QM/MM study and Raman spectroscopy calculations. J.L. performed MD simulations. Y.Z. and L.L. performed the stopped-flow absorption measurements. Z.B. and H.M. assisted in preliminary experiments. B.W., J.-F.L., L.L., Q.-Q.L., X.Z. and J.L. co-wrote the manuscript; B.R., C.W., X.W. and L.Y. critically reviewed the manuscript.

## References

1. Solomon, E. I. et al. Copper active sites in biology. Chem. Rev. 114, 3659–3853 (2014).

2. Tucci, F. J. & Rosenzweig, A. C. Direct methane oxidation by copper- and iron-dependent methane monooxygenases. Chem. Rev. 124, 1288–1320 (2024).

3. Guengerich, F. P. & Yoshimoto, F. K. Formation and cleavage of C–C bonds by enzymatic oxidation–reduction reactions. Chem. Rev. 118, 6573–6655 (2018).

4. Kovaleva, E. G. & Lipscomb, J. D. Versatility of biological non-heme Fe(II) centers in oxygen activation reactions. Nat. Chem. Biol. 4, 186–193 (2008).

5. Islam, M. S., Leissing, T. M., Chowdhury, R., Hopkinson, R. J. & Schofield, C. J. 2-Oxoglutarate-dependent oxygenases. Annu. Rev. Biochem. 87, 585–620 (2018).

6. Jasniewski, A. J. & Que, L., Jr. Dioxygen activation by nonheme diiron enzymes: diverse dioxygen adducts, high-valent intermediates, and related model complexes. Chem. Rev. 118, 2554–2592 (2018).

7. Krebs, C., Galonić Fujimori, D., Walsh, C. T. & Bollinger, J. M., Jr. Non-heme Fe(IV)–oxo intermediates. Acc. Chem. Res. 40, 484–492 (2007).

8. Shaik, S. et al. P450 enzymes: their structure, reactivity, and selectivity—modeled by QM/MM calculations. Chem. Rev. 110, 949–1017 (2010).

9. Huang, X. & Groves, J. T. Oxygen activation and radical transformations in heme proteins and metalloporphyrins. Chem. Rev. 118, 2491–2553 (2018).

10. Ushimaru, R. & Abe, I. Unusual dioxygen-dependent reactions catalyzed by nonheme iron enzymes in natural product biosynthesis. ACS Catal. 13, 1045–1076 (2023).

11. Tang, M.-C., Zou, Y., Watanabe, K., Walsh, C. T. & Tang, Y. Oxidative cyclization in natural product biosynthesis. Chem. Rev. 117, 5226–5333 (2017).

12. Paton, A. E. et al. Connecting chemical and protein sequence space to predict biocatalytic reactions. Nature 646, 108–116 (2025).

13. Lu, H. P., Xun, L. & Xie, X. S. Single-molecule enzymatic dynamics. Science 282, 1877–1882 (1998).

14. Rittle, J. & Green, M. T. Cytochrome P450 compound I: capture, characterization, and C–H bond activation kinetics. Science 330, 933–937 (2010).

15. Matthews, M. L. et al. Substrate-triggered formation and remarkable stability of the C–H bond-cleaving chloroferryl intermediate in the aliphatic halogenase, SyrB2. Biochemistry 48, 4331–4343 (2009).

16. Davydov, R. et al. Hydroxylation of camphor by reduced oxy-cytochrome P450cam: mechanistic implications of EPR and ENDOR studies of catalytic intermediates in native and mutant enzymes. J. Am. Chem. Soc. 123, 1403–1415 (2001).

17. Panay, A. J., Lee, M., Krebs, C., Bollinger, J. M., Jr. & Fitzpatrick, P. F. Evidence for a high-spin Fe(IV) species in the catalytic cycle of a bacterial phenylalanine hydroxylase. Biochemistry 50, 1928–1933 (2011).

18. Neese, F. Quantum chemical calculations of spectroscopic properties of metalloproteins and model compounds: EPR and Mössbauer properties. Curr. Opin. Chem. Biol. 7, 125–135 (2003).

19. Banerjee, R., Proshlyakov, Y., Lipscomb, J. D. & Proshlyakov, D. A. Structure of the key species in the enzymatic oxidation of methane to methanol. Nature 518, 431–434 (2015).

20. Heidinger, L. et al. Analysis of early intermediate states of the nitrogenase reaction by regularization of EPR spectra. Nat. Commun. 15, 4041 (2024).

21. Bollmeyer, M. M. et al. Insights into the initial steps of the thiamin pyrimidine synthase (ThiC)-catalyzed reaction through EPR spectroscopic characterization of radical intermediates. Chem. Sci. 17, 1624–1636 (2026).

22. Balo, A. R., et al. Trapping a cross-linked lysine–tryptophan radical in the catalytic cycle of the radical SAM enzyme SuiB. Proc. Natl. Acad. Sci. USA 118, e2101571118 (2021).

23. Caserta, G. et al. Stepwise assembly of the active site of [NiFe]-hydrogenase. Nat. Chem. Biol. 19, 498–506 (2023).

24. Schulz, V. et al. Mechanism and structural dynamics of sulfur transfer during de novo [2Fe-2S] cluster assembly on ISCU2. Nat. Commun. 15, 3269 (2024).

25. Iyer, S. R., et al. Direct coordination of pterin to Fe^II^ enables neurotransmitter biosynthesis in the pterin-dependent hydroxylases. Proc. Natl. Acad. Sci. USA 118, e2022379118 (2021).

26. Satpathy, J. K. et al. Mechanistic divergence in sulfur-ligated iron(III)-alkylperoxo reactivity: aldehyde oxidation prevails over deformylation. Angew. Chem. Int. Ed. 64, e202512839 (2025).

27. Rousseau, D. L., Ishigami, I. & Yeh, S.-R. Structural and functional mechanisms of cytochrome c oxidase. J. Inorg. Biochem. 262, 112730 (2025).

28. Kipouros, I., et al. Elucidation of the tyrosinase/O2/monophenol ternary intermediate that dictates the monooxygenation mechanism in melanin biosynthesis. Proc. Natl. Acad. Sci. USA 119, e2205619119 (2022).

29. Srinivas, V. et al. High-resolution XFEL structure of the soluble methane monooxygenase hydroxylase complex with its regulatory component at ambient temperature in two oxidation states. J. Am. Chem. Soc. 142, 14249–14266 (2020).

30. Ohmer, C. J. et al. XFEL serial crystallography reveals the room temperature structure of methyl-coenzyme M reductase. J. Inorg. Biochem. 230, 111768 (2022).

31. Proshlyakov, D. A., McCracken, J. & Hausinger, R. P. Spectroscopic analyses of 2-oxoglutarate-dependent oxygenases: TauD as a case study. J. Biol. Inorg. Chem. 22, 367–379 (2017).

32. Zhang, A. et al. Catalytic cycle of formate dehydrogenase captured by single-molecule conductance. Nat. Catal. 6, 266–275 (2023).

33. Chen, Y., Huang, J., Wu, S., Zhang, A. & Hong, W. Single-molecule protein studies using scanning tunneling microscope break junction technique. Trends Chem. 8, 459–473 (2026).

34. Hess, C. New advances in using Raman spectroscopy for the characterization of catalysts and catalytic reactions. Chem. Soc. Rev. 50, 3519–3564 (2021).

35. Buhrke, D. & Hildebrandt, P. Probing structure and reaction dynamics of proteins using time-resolved resonance Raman spectroscopy. Chem. Rev. 120, 3577–3630 (2020).

36. Haynes, C. L., McFarland, A. D. & Van Duyne, R. P. Surface-enhanced Raman spectroscopy. Anal. Chem. 77, 338 A–346 A (2005).

37. Yi, J. et al. Surface-enhanced Raman spectroscopy: a half-century historical perspective. Chem. Soc. Rev. 54, 1453–1551 (2025).

38. Langer, J. et al. Present and future of surface-enhanced Raman scattering. ACS Nano 14, 28–117 (2020).

39. Li, J.-F. & Tian, Z.-Q. Shell-isolated nanoparticle-enhanced Raman spectroscopy (SHINERS), in Frontiers of surface-enhanced Raman scattering: single nanoparticles and single cells, ed. by Y. Ozaki, K. Kneipp, R. Aroca (John Wiley & Sons, Chichester, 2014), pp. 163–192 (2014).

40. Ding, S.-Y. et al. Nanostructure-based plasmon-enhanced Raman spectroscopy for surface analysis of materials. Nat. Rev. Mater. 1, 16021 (2016).

41. Zong, C. et al. Surface-enhanced Raman spectroscopy for bioanalysis: reliability and challenges. Chem. Rev. 118, 4946–4980 (2018).

42. Li, J. F. et al. Shell-isolated nanoparticle-enhanced Raman spectroscopy. Nature 464, 392–395 (2010).

43. Zhang, H. et al. In situ dynamic tracking of heterogeneous nanocatalytic processes by shell-isolated nanoparticle-enhanced Raman spectroscopy. Nat. Commun. 8, 15447 (2017).

44. Pérez-Jiménez, A. I., Lyu, D., Lu, Z., Liu, G. & Ren, B. Surface-enhanced Raman spectroscopy: benefits, trade-offs and future developments. Chem. Sci. 11, 4563–4577 (2020).

45. An, H., et al. Sub-second time-resolved surface-enhanced Raman spectroscopy reveals dynamic CO intermediates during electrochemical CO_2_ reduction on copper. Angew. Chem. Int. Ed. 60, 16576–16584 (2021).

46. Zhang, Y.-J. et al. Shell-isolated nanoparticle-enhanced Raman spectroscopy. Nat. Rev. Methods Primers 3, 36 (2023).

47. Dong, J.-C. et al. In situ Raman spectroscopic evidence for oxygen reduction reaction intermediates at platinum single-crystal surfaces. Nat. Energy 4, 60–67 (2019).

48. Price, J. C., Barr, E. W., Tirupati, B., Bollinger, J. M. & Krebs, C. The first direct characterization of a high-valent iron intermediate in the reaction of an α-ketoglutarate-dependent dioxygenase: a high-spin Fe(IV) complex in taurine/α-ketoglutarate dioxygenase (TauD) from *Escherichia coli*. Biochemistry 42, 7497–7508 (2003).

49. Grzyska, P. K., Müller, T. A., Campbell, M. G. & Hausinger, R. P. Metal ligand substitution and evidence for quinone formation in taurine/α-ketoglutarate dioxygenase. J. Inorg. Biochem. 101, 797–808 (2007).

50. Sinnecker, S. et al. Spectroscopic and computational evaluation of the structure of the high-spin Fe(IV)-oxo intermediates in taurine: α-ketoglutarate dioxygenase from *Escherichia coli* and its His99Ala ligand variant. J. Am. Chem. Soc. 129, 6168–6179 (2007).

51. Proshlyakov, D. A., Henshaw, T. F., Monterosso, G. R., Ryle, M. J. & Hausinger, R. P. Direct detection of oxygen intermediates in the non-heme Fe enzyme taurine/α-ketoglutarate dioxygenase. J. Am. Chem. Soc. 126, 1022–1023 (2004).

52. Mitchell, A. J. et al. Visualizing the reaction cycle in an iron(II)- and 2-(oxo)-glutarate-dependent hydroxylase. J. Am. Chem. Soc. 139, 13830–13836 (2017).

53. Kellner, D. G., Hung, S.-C., Weiss, K. E. & Sligar, S. G. Kinetic characterization of compound I formation in the thermostable cytochrome P450 CYP119. J. Biol. Chem. 277, 9641–9644 (2002).

54. Terner, J. et al. Resonance Raman spectroscopy of oxoiron(IV) porphyrin π-cation radical and oxoiron(IV) hemes in peroxidase intermediates. J. Inorg. Biochem. 100, 480–501 (2006).

55. Bunno, R., Awakawa, T., Mori, T. & Abe, I. Aziridine formation by a Fe^II^/α-ketoglutarate dependent oxygenase and 2-aminoisobutyrate biosynthesis in fungi. Angew. Chem. Int. Ed. 60, 15827–15831 (2021).

56. Tao, H. et al. Stereoselectivity and substrate specificity of the Fe(II)/α-ketoglutarate-dependent oxygenase TqaL. J. Am. Chem. Soc. 144, 21512–21520 (2022).

57. Zhang, X., Liu, J., Liao, L., Wang, Z. & Wang, B. Coordination dynamics of iron enables the selective C–N coupling but bypasses unwanted C–H hydroxylation in Fe(II)/α-ketoglutarate-dependent non-heme enzymes. Chin. J. Catal. 62, 131–144 (2024).

58. Kissman, E. N. et al. Dynamic metal coordination controls chemoselectivity in a radical halogenase. Nat. Chem. Biol. 22, 491–500 (2026).

59. Palacios, P. M. et al. Regio- and stereoselective halogenation by an iron(II)- and 2-oxoglutarate-dependent halogenase in the biosynthesis of halogenated nucleosides. J. Am. Chem. Soc. 147, 48166–48179 (2025).

60. Heim, P. et al. Aromatic and aliphatic hydrocarbon hydroxylation via a formally Ni^IV^═O oxidant. Dalton Trans. 52, 2663–2671 (2023).

61. Frens, G. Controlled nucleation for the regulation of the particle size in monodisperse gold suspensions. Nat. Phys. Sci. 241, 20–22 (1973).

62. Liao, L. et al. Construction and characterization of a novel glucose dehydrogenase-leucine dehydrogenase fusion enzyme for the biosynthesis of l-tert-leucine. Microb. Cell Fact. 20, 3 (2021).

63. Aslantas, Y. & Surmeli, N. B. Effects of N-terminal and C-terminal polyhistidine tag on the stability and function of the thermophilic P450 CYP119. Bioinorg. Chem. Appl. 2019, 8080697 (2019).

64. Eichhorn, E., van der Ploeg, J. R., Kertesz, M. A. & Leisinger, T. Characterization of α-ketoglutarate-dependent taurine dioxygenase from *Escherichia coli*. J. Biol. Chem. 272, 23031–23036 (1997).

65. Yin, X. & Zabriskie, T. M. VioC is a non-heme iron, α-ketoglutarate-dependent oxygenase that catalyzes the formation of 3S-hydroxy-L-arginine during viomycin biosynthesis. ChemBioChem 5, 1274–1277 (2004).

66. Liao, L. et al. A molecular mechanism for the reduction of the Cu site in lytic polysaccharide monooxygenases by phenol reductants. JACS Au 5, 4233–4248 (2025).

67. Berry, E. A. & Trumpower, B. L. Simultaneous determination of hemes a, b, and c from pyridine hemochrome spectra. Anal. Biochem. 161, 1–15 (1987).

68. Lim, Y. R. et al. Regioselective oxidation of lauric acid by CYP119, an orphan cytochrome P450 from *Sulfolobus acidocaldarius*. J. Microbiol. Biotechnol. 20, 574–578 (2010).

69. Durst, H. D., Milano, M., Kikta, E. J., Connelly, S. A. & Grushka, E. Phenacyl esters of fatty acids via crown ether catalysts for enhanced ultraviolet detection in liquid chromatography. Anal. Chem. 47, 1797–1801 (1975).

70. Aoyama, T. & Sato, R. High pressure liquid chromatography methods for separation of *ω*- and (*ω*-1)-hydroxy fatty acids: Their applications to microsomal fatty acid *ω*-oxidation. Anal. Biochem. 170, 73–82 (1988).

71. Li, P. & Merz, K. M. Jr. MCPB.py: A python based metal center parameter builder. J. Chem. Inf. Model. 56, 599–604 (2016).

72. Heinz, H., Lin, T. J., Mishra, R. K. & Emami, F. S. Thermodynamically consistent force fields for the assembly of inorganic, organic, and biological nanostructures: The INTERFACE force field. Langmuir 29, 1754–1765 (2013).

73. Heinz, H. The role of chemistry and pH of solid surfaces for specific adsorption of biomolecules in solution—accurate computational models and experiment. J. Phys.: Condens. Matter 26, 244105 (2014).

74. Emami, F. S. et al. Force field and a surface model database for silica to simulate interfacial properties in atomic resolution. Chem. Mater. 26, 2647–2658 (2014).

75. Martínez, L., Andrade, R., Birgin, E. G. & Martínez, J. M. PACKMOL: A package for building initial configurations for molecular dynamics simulations. J. Comput. Chem. 30, 2157–2164 (2009).

76. Sherwood, P., et al. QUASI: A general purpose implementation of the QM/MM approach and its application to problems in catalysis. J. Mol. Struct.: THEOCHEM 632, 1–28 (2003).

77. Smith, W. & Forester, T. R. DL_POLY_2.0: A general-purpose parallel molecular dynamics simulation package. J. Mol. Graphics. 14, 136–141 (1996).

78. Bakowies, D. & Thiel, W. Hybrid models for combined quantum mechanical and molecular mechanical approaches. J. Phys. Chem. 100, 10580–10594 (1996).

