## Supplementary Information for "Ambient-Condition Visualization of Metalloenzyme Reactive Intermediates Enabled by the SHINERS Paradigm"

**This file includes:**

Supplementary Figs. 1-34

Supplementary Table 1

Amino acid sequences 1-4

### Supplementary Figs

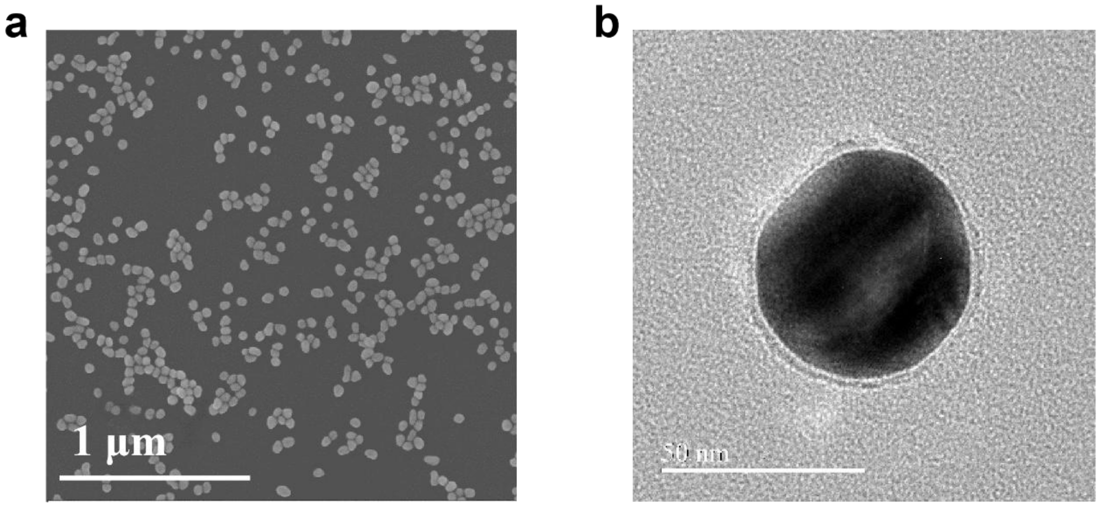


**Supplementary Fig. 1 | Structural characterization of the shell-isolated nanoparticles.** a, Scanning electron microscopy image of the Au@$SiO_2$ nanoparticles. b, High-resolution transmission electron microscopy image of an individual Au@$SiO_2$ nanoparticle, showing the Au core and surrounding silica shell.

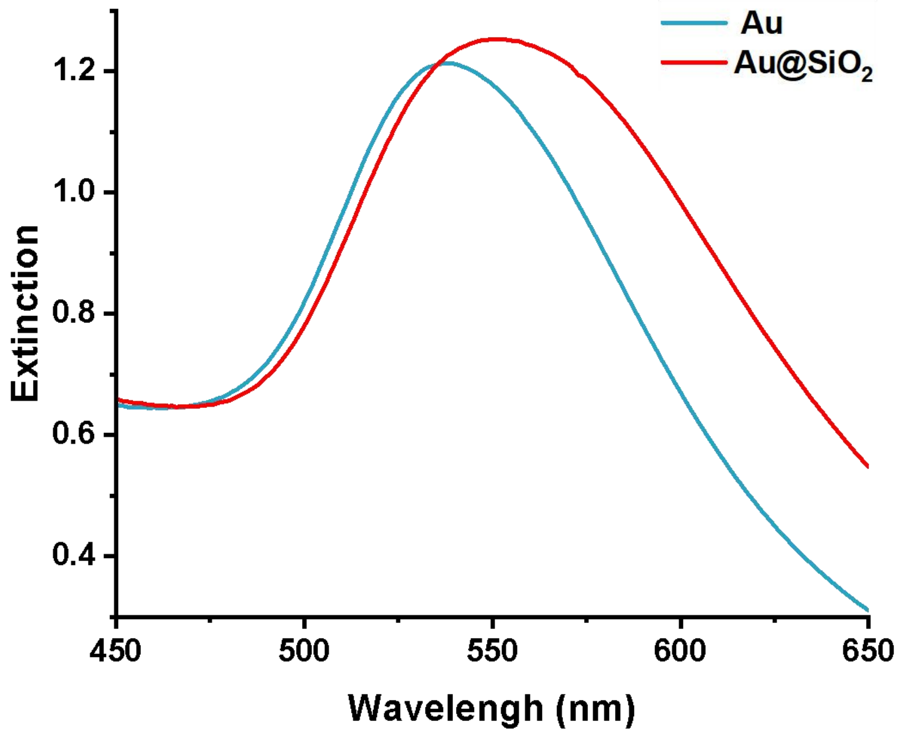


**Supplementary Fig. 2 | Optical characterization of the Au and Au@SiO$_2$ nanoparticles used for SHINERS measurements.** UV–visible extinction spectra of the bare Au nanoparticles and the corresponding Au@SiO$_2$ shell-isolated nanoparticles. The localized surface plasmon resonance (LSPR) extinction maximum shifts from approximately 540 nm for Au to 552 nm for Au@SiO$_2$ after formation of the ultrathin SiO$_2$ shell, indicating retention of the characteristic plasmonic response of the Au core after silica coating.

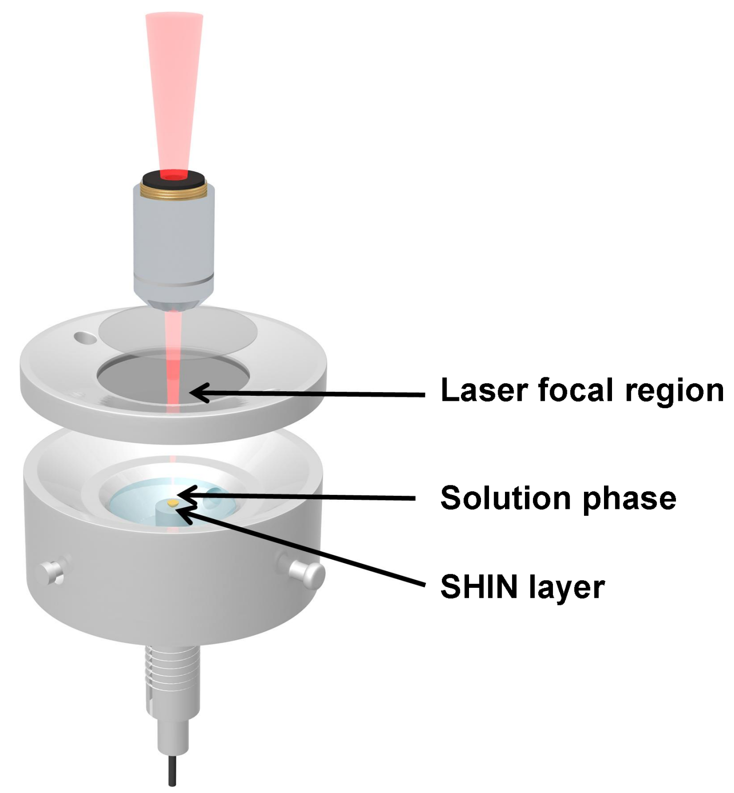


**Supplementary Fig. 3 | Raman cell used for interfacial SHINERS measurements.** Schematic of the homemade Raman cell showing the SHIN layer, aqueous sample phase, and laser focal region at the solid/liquid interface.

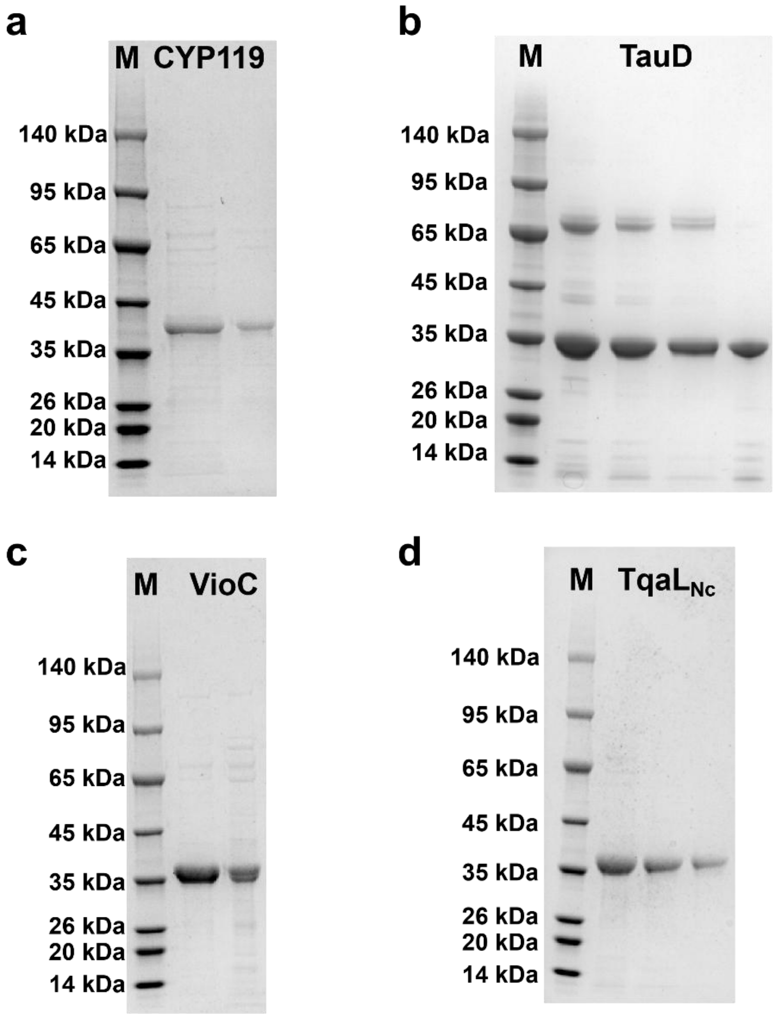


**Supplementary Fig. 4 | SDS-PAGE analysis of the purified metalloenzymes.** a, CYP119. b, TauD. c, VioC. d, TqaLNc. Molecular-mass markers are indicated by M.

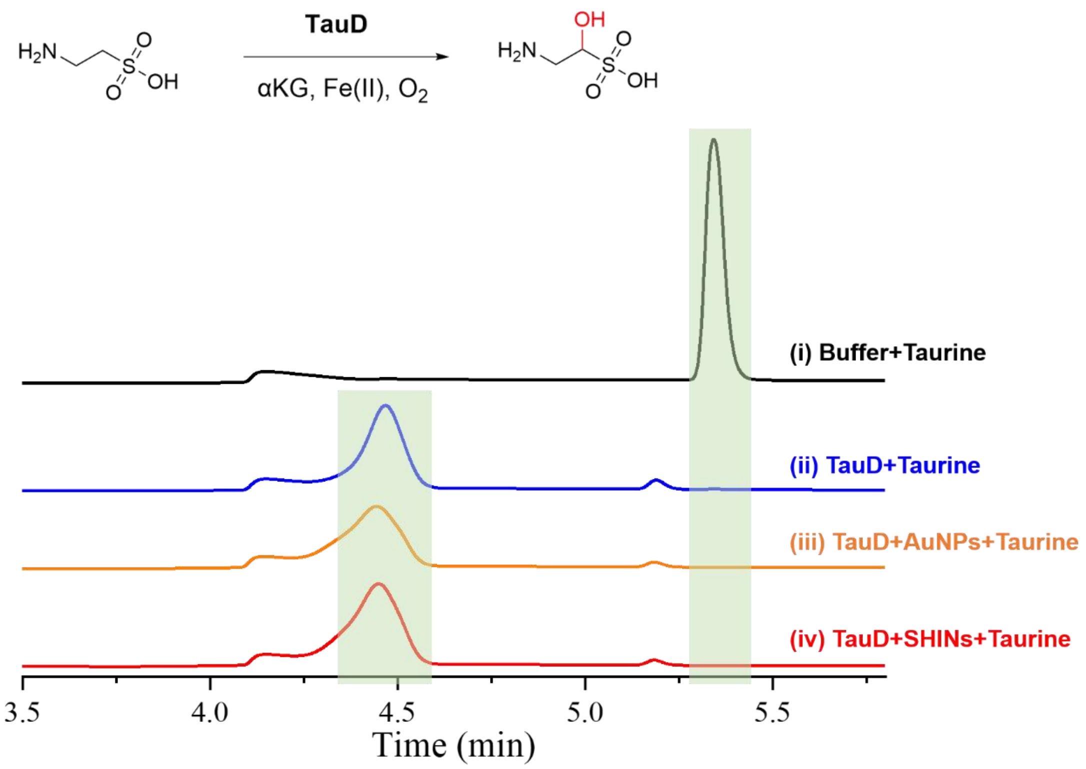


**Supplementary Fig. 5 | Effect of nanoparticles on TauD-catalyzed taurine hydroxylation.** HPLC chromatograms for the blank control containing buffer in place of TauD (i), the complete TauD reaction (ii), and complete reactions after pre-incubation with AuNPs (iii) or SHINs (iv). Standard conditions were 20 µM TauD, 1 mM taurine, 2 mM αKG, and 200 µM $FeSO_4$ in 150 mM Tris-HCl (pH 8.0) containing 300 mM NaCl at 30 °C. This measurement serves as a qualitative catalytic-competence control for the presence of nanoparticles and is not used for quantitative comparison of catalytic rates or for spectroscopic assignment of the SHINERS features.

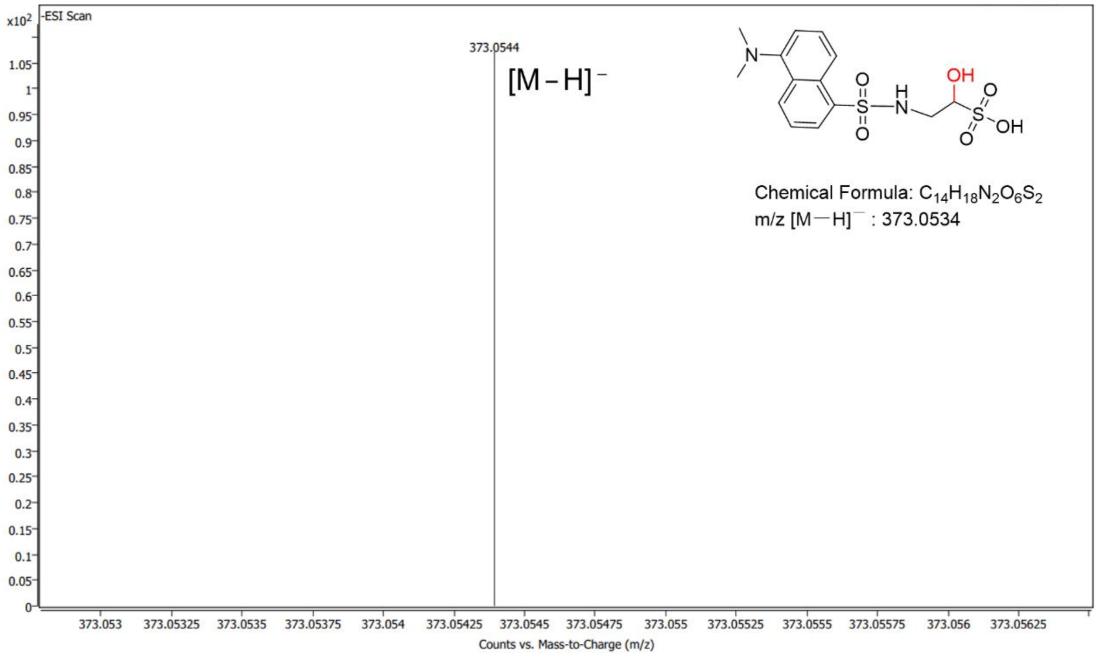


**Supplementary Fig. 6 | LC-MS identification of the TauD hydroxylation product.** The TauD reaction with taurine was performed at 30 °C for 1 h. The product was derivatized with dansyl chloride and analyzed by LC-MS. The observed ion is consistent with the dansylated hydroxylated product.

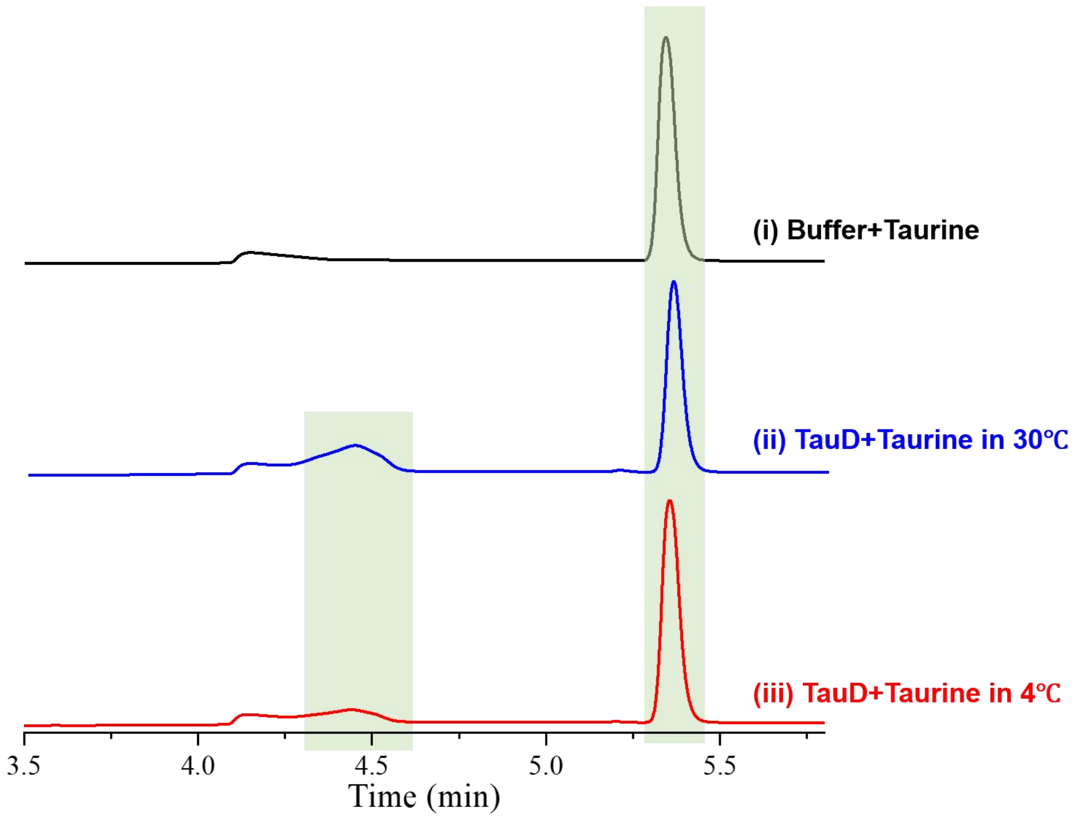


**Supplementary Fig. 7 | Temperature dependence of TauD-catalyzed taurine hydroxylation.** HPLC chromatograms for the blank control at 30 °C (i), the complete reaction at 30 °C (ii), and the complete reaction at 4 °C (iii). Reaction mixtures contained 20 μM TauD, 1 mM taurine, 2 mM αKG, and 200 μM $FeSO_4$ in 150 mM Tris-HCl (pH 8.0) containing 300 mM NaCl. This experiment confirms that TauD product formation remains detectable at 4 °C, the temperature used for the SHINERS measurements, and is not intended to provide temperature-dependent kinetic parameters.

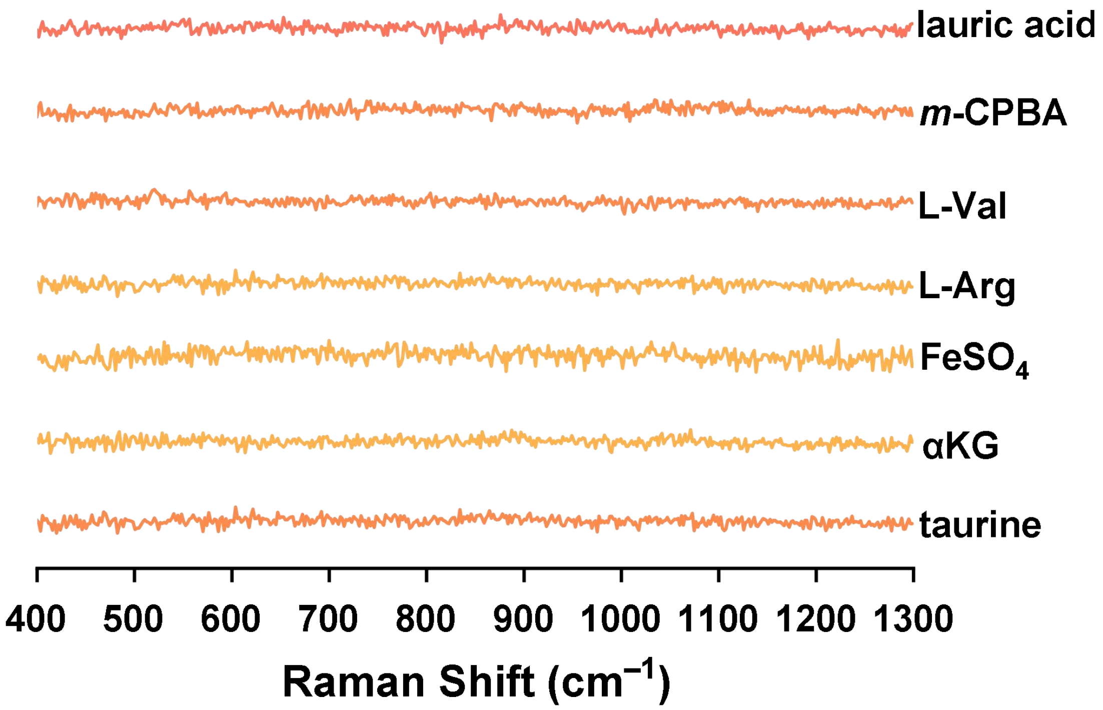


**Supplementary Fig. 8 | Raman spectra of individual reaction components used to define spectral backgrounds.** Spectra were recorded separately for 1 mM taurine, 2 mM αKG, 200 µM $FeSO_4$, 1 mM L-arginine, 1 mM L-valine, 1 mM *m*-CPBA, and 1 mM lauric acid as reference spectra using the same Raman measurement setup. These spectra serve as reference backgrounds for evaluating reaction-associated spectral changes; the presence or absence of an individual component band is not, by itself, used to assign a reactive intermediate.

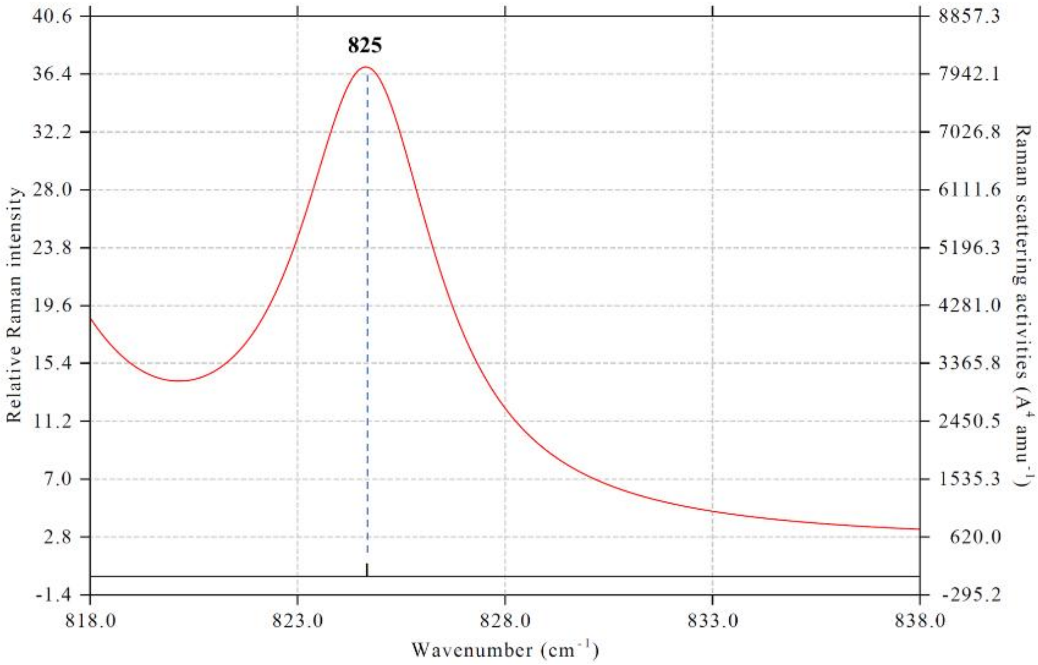


**Supplementary Fig. 9 | Harmonic QM/MM Raman mode calculated for the TauD ferryl model.** Calculated Raman activity for the standard-isotope QM/MM-optimized Fe(IV)=O model, showing the Fe–O stretching mode near 825 cm$^{-1}$. No anharmonic correction or empirical frequency-scaling factor was applied. The calculation is used primarily to evaluate Fe–O vibrational mode character and the approximate unscaled frequency; the experimental oxygen-isotope shift is evaluated independently and is not derived from this standard-isotope calculation.

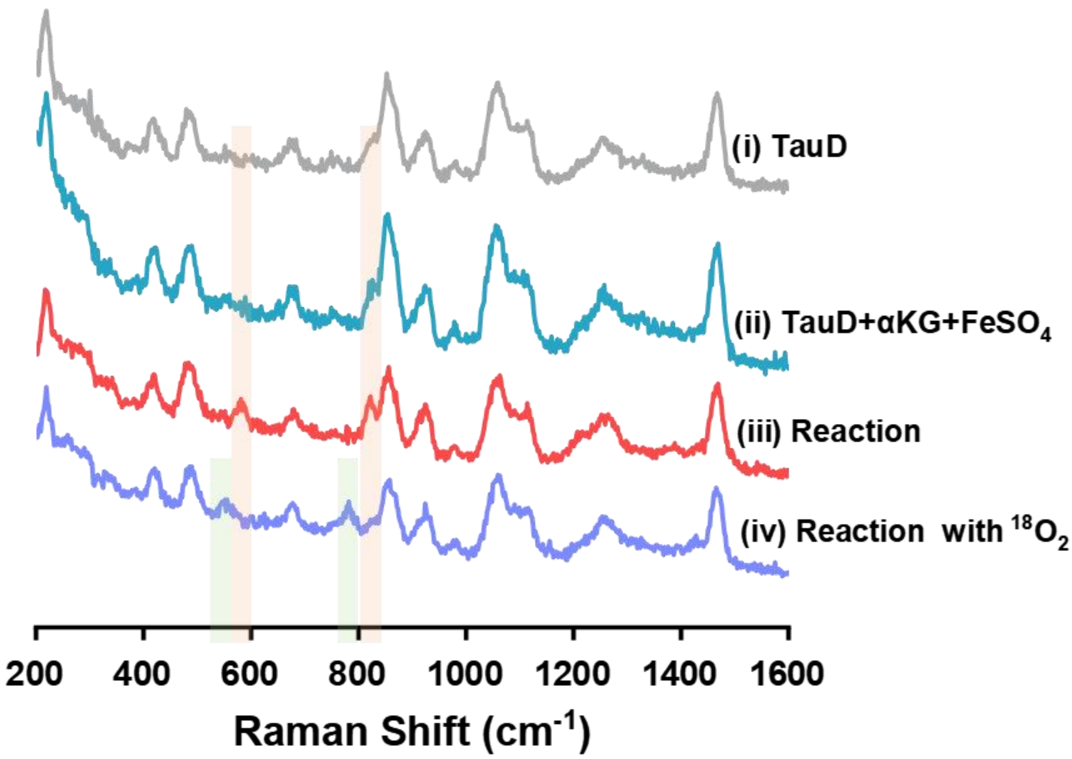


**Supplementary Fig. 10 | Unprocessed raw SHINERS spectra of the TauD reaction.** Spectra shown without baseline correction for i, TauD alone; ii, the pre-reaction mixture containing TauD, αKG, and $FeSO_4$; iii, the complete reaction after taurine addition; and iv, the corresponding reaction under an $^{18}O_2$ atmosphere. The shaded region denotes the spectral region containing the ferryl-associated feature discussed in the main text and is provided only as a visual guide. These unprocessed spectra are provided to permit direct assessment of the spectral background and isotope-dependent peak positions independently of baseline correction.

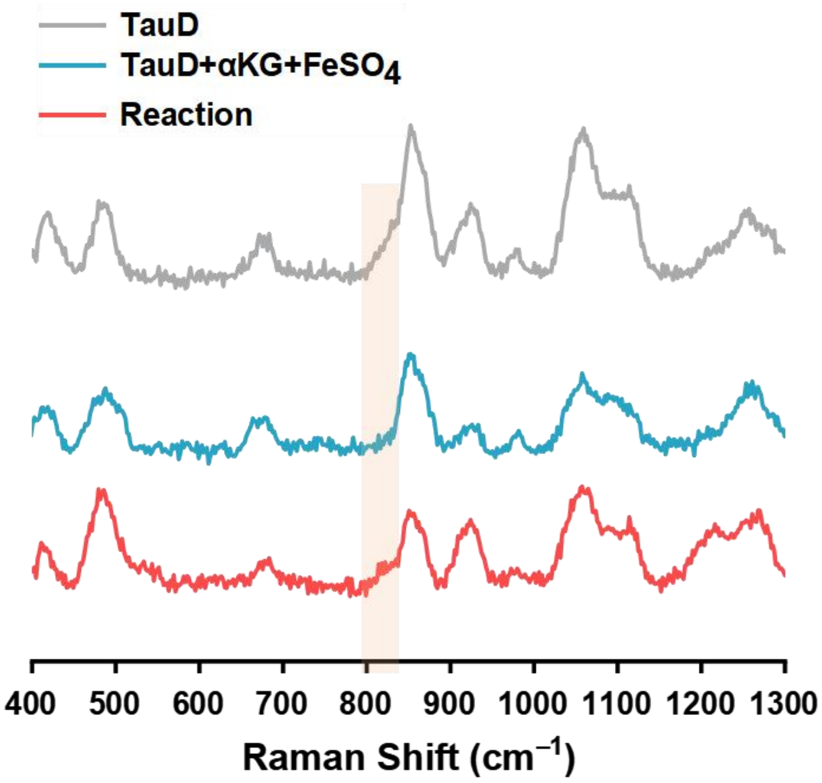


**Supplementary Fig. 11 | TauD Raman spectra obtained with silica-uncoated Au nanoparticles.** SERS spectra of TauD alone, the pre-reaction mixture containing TauD, αKG, and $FeSO_4$, and the complete reaction after taurine addition, acquired using silica-uncoated Au nanoparticles in place of Au@$SiO_2$ SHINs. The shaded region denotes the expected ferryl spectral region and does not indicate an independently resolved Fe(IV)=O band under the bare-Au condition. This control is used qualitatively to assess the consequence of removing the silica isolation layer; because direct biomolecule–Au interactions can alter adsorption and the Raman background, it is not used for quantitative comparison of enhancement factors.

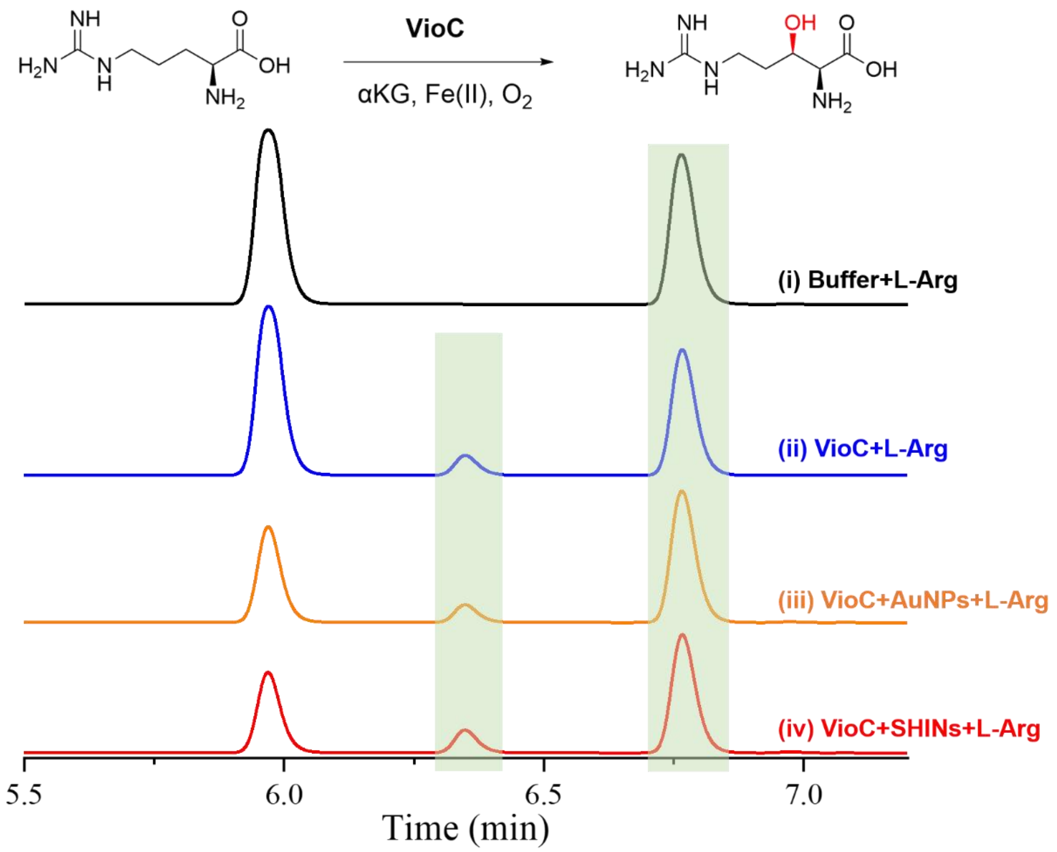


**Supplementary Fig. 12 | Effect of nanoparticles on VioC-catalyzed L-arginine hydroxylation.** HPLC chromatograms for the blank control containing buffer in place of VioC (i), the complete VioC reaction (ii), and complete reactions after pre-incubation with AuNPs (iii) or SHINs (iv). Standard conditions were 20 μM VioC, 1 mM L-arginine, 2 mM αKG, and 200 μM $FeSO_4$ in 150 mM Tris-HCl (pH 8.0) containing 300 mM NaCl at 30 °C. This measurement serves as a qualitative catalytic-competence control for the presence of nanoparticles and is not used for quantitative comparison of catalytic rates or for assignment of the approximately 816 $cm^{-1}$ Raman feature.

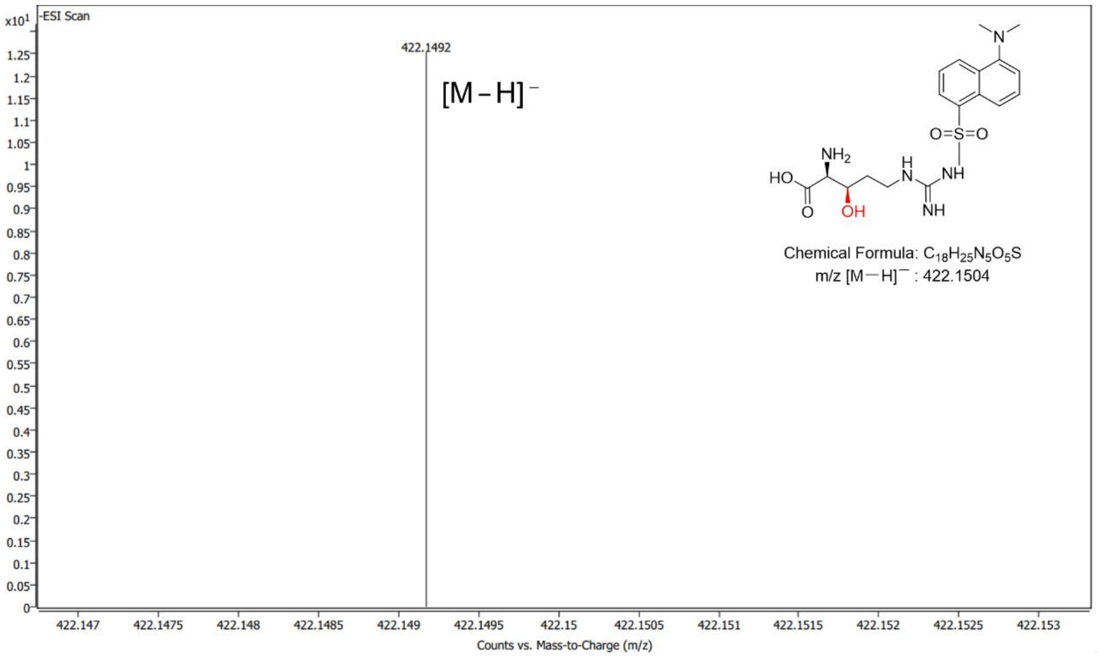


**Supplementary Fig. 13 | LC-MS identification of the VioC hydroxylation product.** The VioC reaction with L-arginine was performed at 30 °C for 1 h. The product was derivatized with dansyl chloride and analyzed by LC-MS. The observed ion is consistent with the dansylated hydroxylated product.

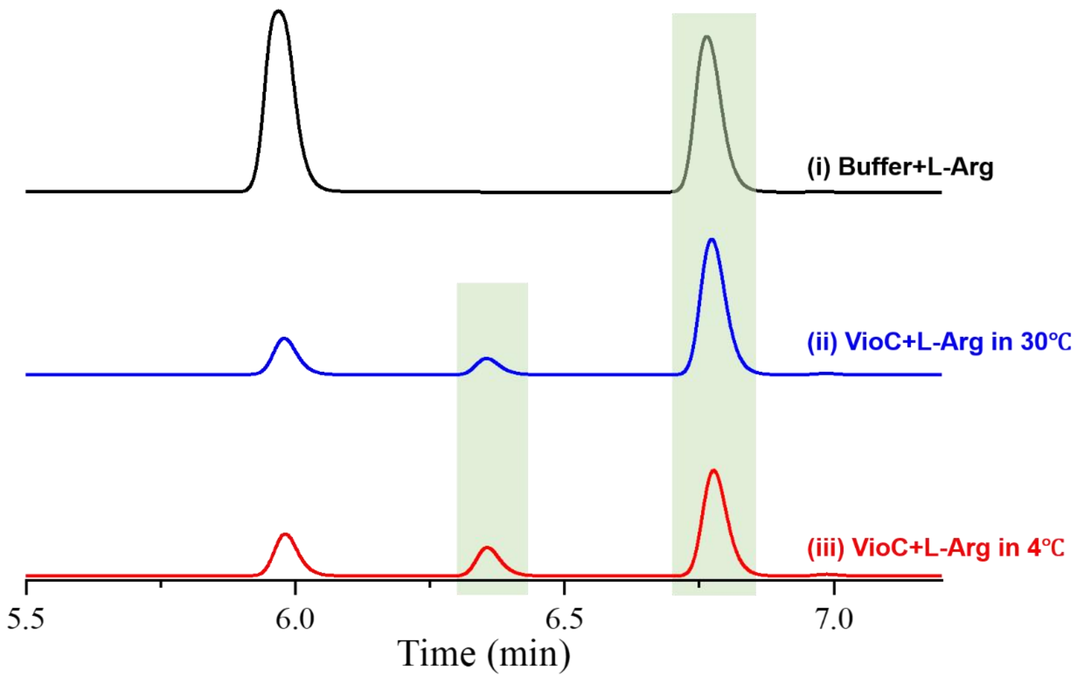


**Supplementary Fig. 14 | Temperature dependence of VioC-catalyzed L-arginine hydroxylation.** HPLC chromatograms for the blank control at 30 °C (i), the complete reaction at 30 °C (ii), and the complete reaction at 4 °C (iii). Reaction mixtures contained 20 µM VioC, 1 mM L-arginine, 2 mM αKG, and 200 µM $FeSO_4$ in 150 mM Tris-HCl (pH 8.0) containing 300 mM NaCl. This experiment confirms that VioC product formation remains detectable at 4 °C, the temperature used for the SHINERS measurements, and is not intended to provide temperature-dependent kinetic parameters.

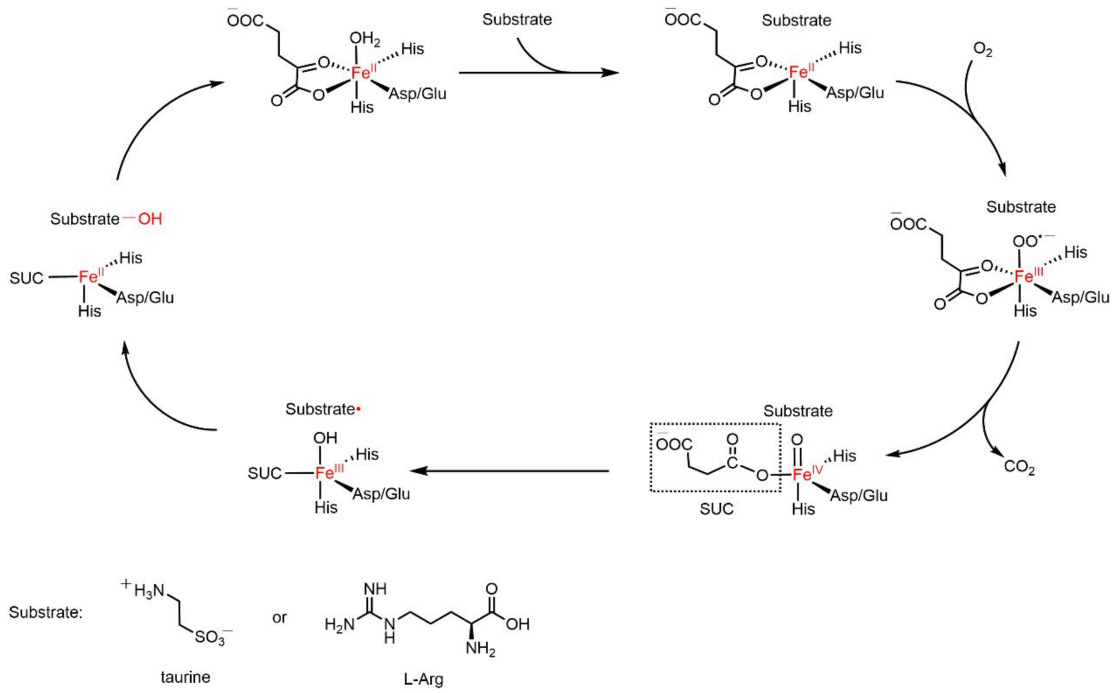


**Supplementary Fig. 15 | Proposed catalytic cycle of Fe(II)/α-ketoglutarate-dependent hydroxylases.** The scheme summarizes oxygen activation, formation of an Fe(IV)=O state, substrate C–H activation, hydroxyl rebound, and product release for hydroxylation chemistry in TauD and VioC. This scheme is shown for mechanistic context and does not imply that every intermediate or individual step in the catalytic cycle was directly observed by SHINERS in the present study.

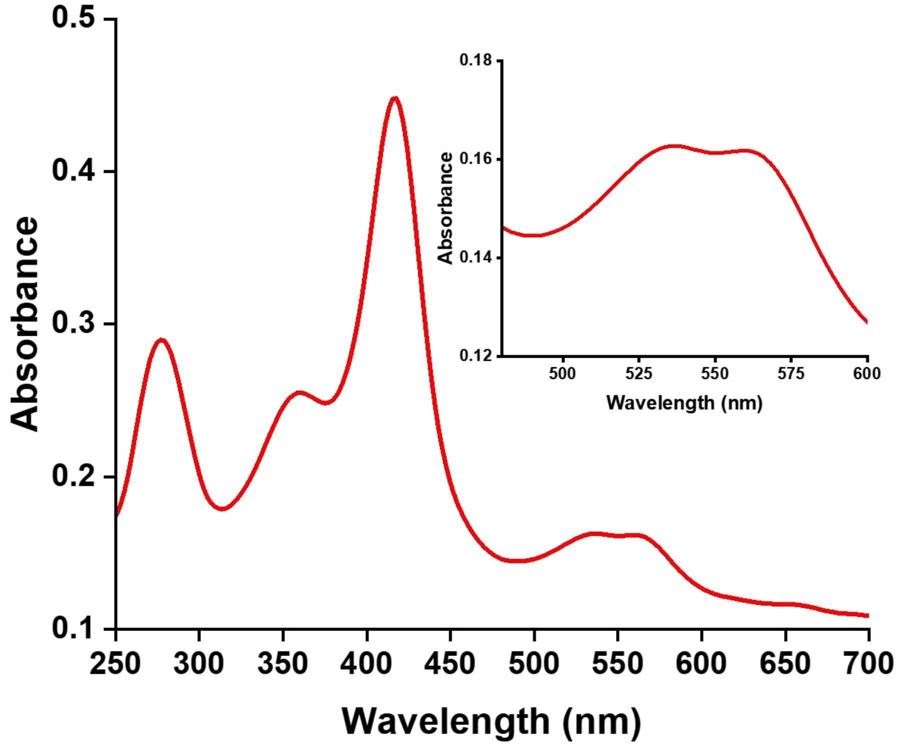


**Supplementary Fig. 16 | UV–visible absorption spectrum of purified ferric CYP119.** UV–visible absorption spectrum of freshly prepared ferric CYP119 recorded before the stopped-flow and SHINERS measurements. The spectrum displays the characteristic Soret and Q-band features of the resting ferric enzyme. The inset shows an expanded view of the Q-band region.

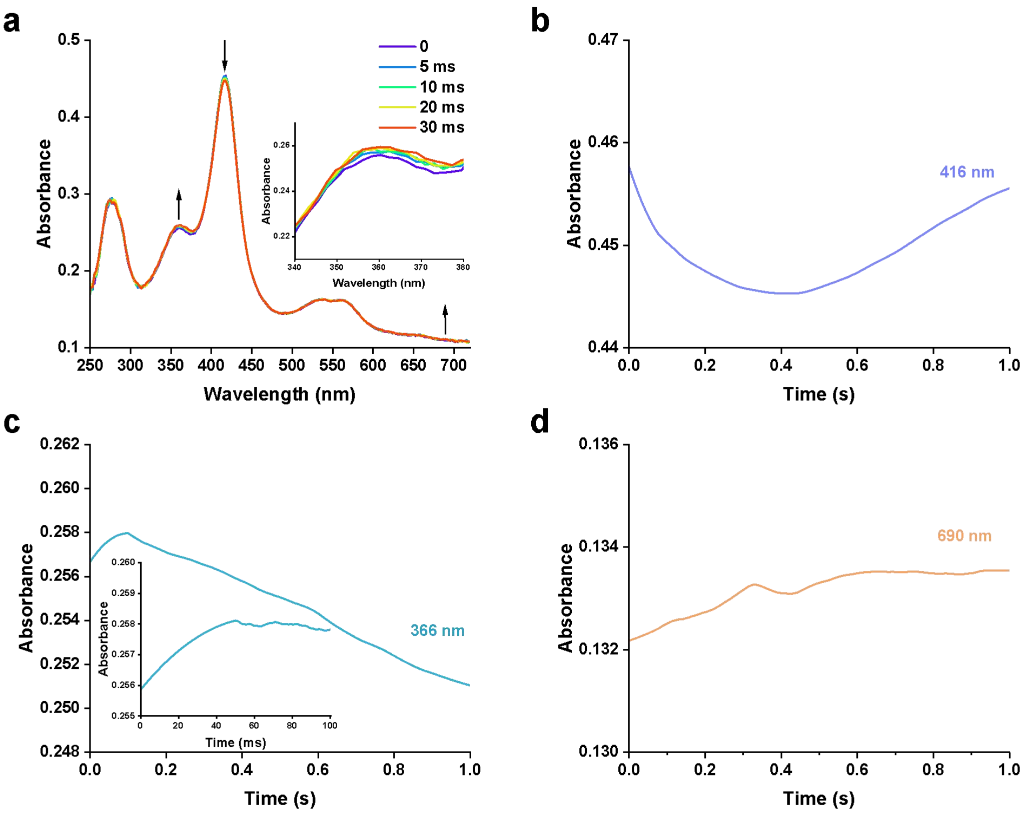


**Supplementary Fig. 17 | Early-time stopped-flow absorption spectra of the CYP119/*m*-CPBA reaction.** CYP119 was rapidly mixed with *m*-CPBA to give final concentrations of 10 µM CYP119 and 20 µM *m*-CPBA in 100 mM potassium phosphate buffer (pH 7.0) at 4 °C. **a**, Early-time absorption spectra collected after rapid mixing; representative spectra at 0, 5, 10, 20, and 30 ms are shown. Arrows indicate the principal directions of the early spectral changes. **b–d**, Corresponding kinetic traces monitored at 416, 366, and 690 nm, respectively. These measurements are used to characterize the early-time electronic spectral evolution of the CYP119/*m*-CPBA reaction and are not taken as a stand-alone spectroscopic identification of Compound I or as evidence for its persistence during the 120 s SHINERS acquisition.

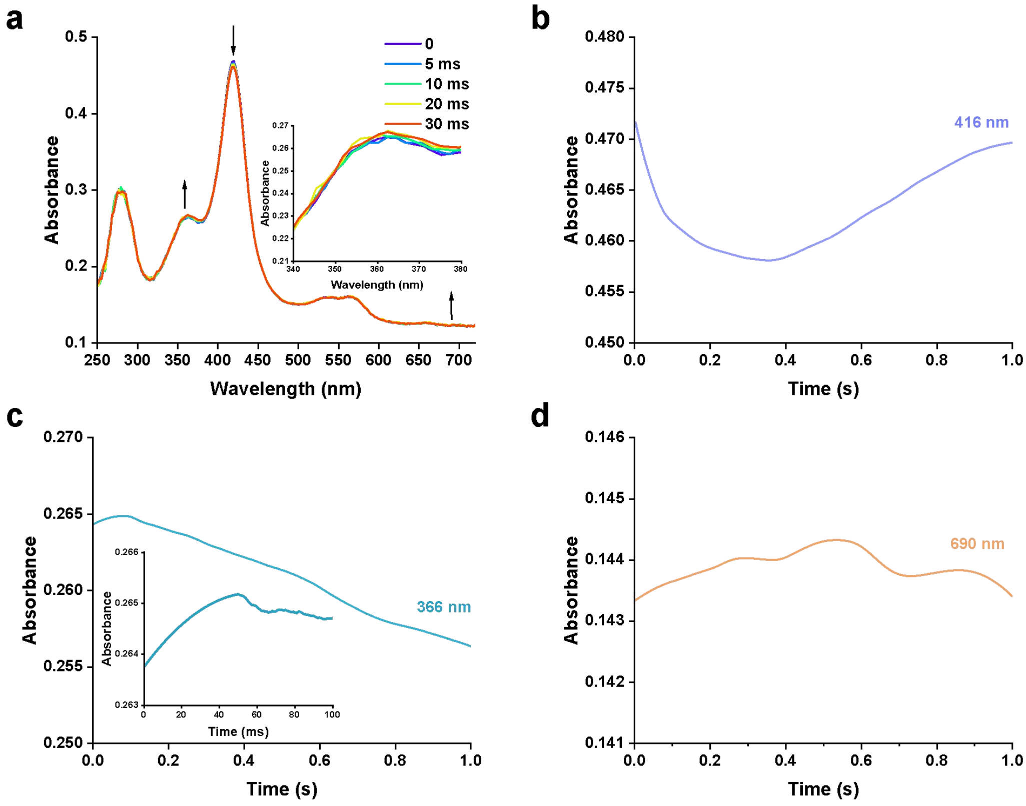


**Supplementary Fig. 18 | Effect of SHINs on early-time CYP119/*m*-CPBA spectral evolution.** Matched stopped-flow measurements were performed by rapidly mixing CYP119 and *m*-CPBA in the presence of SHINs to give final concentrations of 10 μM CYP119 and 20 μM *m*-CPBA at 4 °C. **a**, Early-time absorption spectra collected after rapid mixing; representative spectra at 0, 5, 10, 20, and 30 ms are shown. **b–d**, Corresponding kinetic traces monitored at 416, 366, and 690 nm, respectively. These matched measurements are used to assess whether the presence of SHINs substantially perturbs the early-time electronic spectral evolution of the CYP119/*m*-CPBA system; they are not used to determine a Compound I lifetime or to infer the kinetics of the time-integrated SHINERS signal.

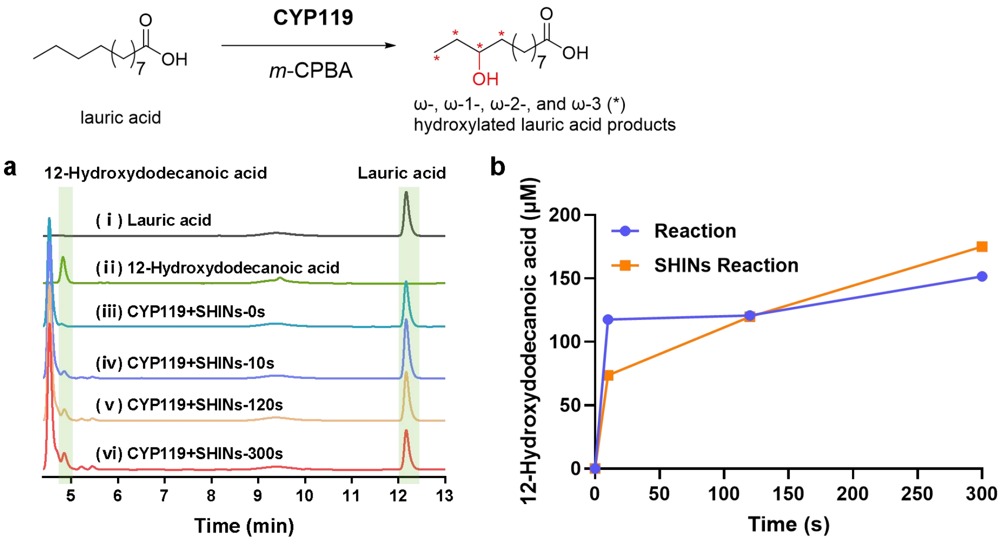


**Supplementary Fig. 19 | CYP119 catalytic competence in the presence of SHINs.** Time-dependent product formation during CYP119-catalyzed hydroxylation of lauric acid by *m*-CPBA. Reaction conditions: 20 μM CYP119, 1 mM m-CPBA, 1 mM lauric acid in 100 mM potassium phosphate buffer (pH 7.0) at 4 °C. **a**, Representative HPLC chromatograms of lauric acid, authentic 12-hydroxydodecanoic acid, and CYP119 reaction samples collected at the indicated times in the presence of SHINs. **b**, Time-dependent formation of 12-hydroxydodecanoic acid in matched CYP119 reactions performed with and without SHINs. The hydroxylated product was quantified against an authentic 12-hydroxydodecanoic acid standard. This experiment serves solely as a functional control demonstrating retention of CYP119 catalytic competence in the presence of SHINs. Product formation is not used as a stand-alone basis for assigning the ~828 $cm^{-1}$ Raman feature and is not used to infer the lifetime or instantaneous population of Compound I.

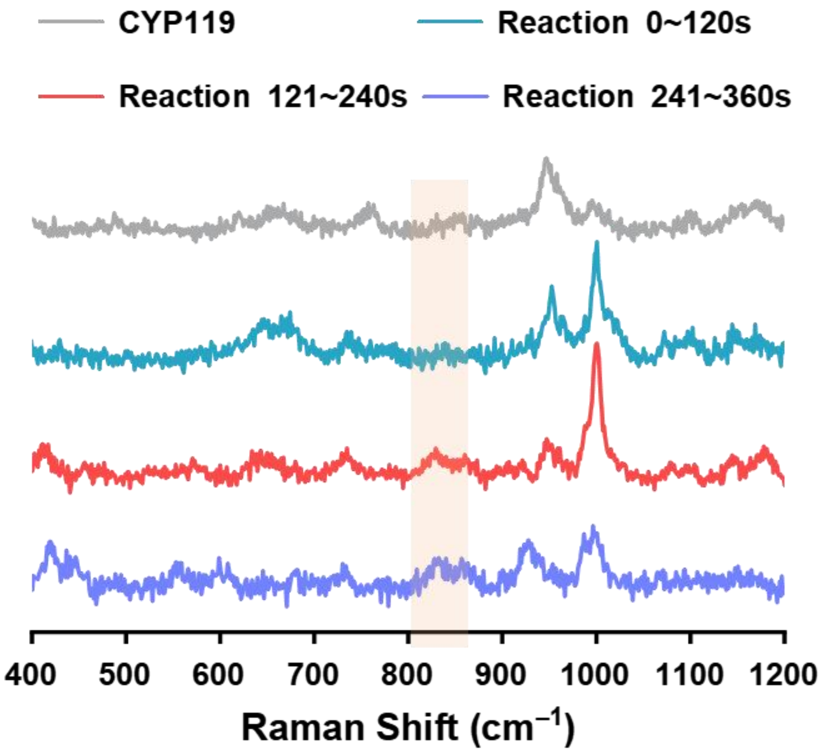


**Supplementary Fig. 20 | Sequential SHINERS acquisition following initiation of the CYP119/*m*-CPBA reaction.** In this timing-control experiment, Raman acquisition was initiated before *m*-CPBA was introduced directly into the Raman cell, ensuring that reaction initiation occurred within the first acquisition window. Consecutive spectra were collected over 0–120, 120–240, and 240–360 s after reaction initiation. The shaded region denotes the spectral region containing the ~828 cm$^{-1}$ Fe(IV)=O feature assigned to CYP119 Compound I in the main text. The ~828 cm$^{-1}$ response is observed across consecutive acquisition windows, consistent with accumulation of Raman photons from the reacting CYP119/*m*-CPBA ensemble. These sequential spectra characterize the evolution of the accumulated interfacial SHINERS signal and are not interpreted as an intrinsic solution-phase kinetic trace or as a measurement of the lifetime or instantaneous population of Compound I.

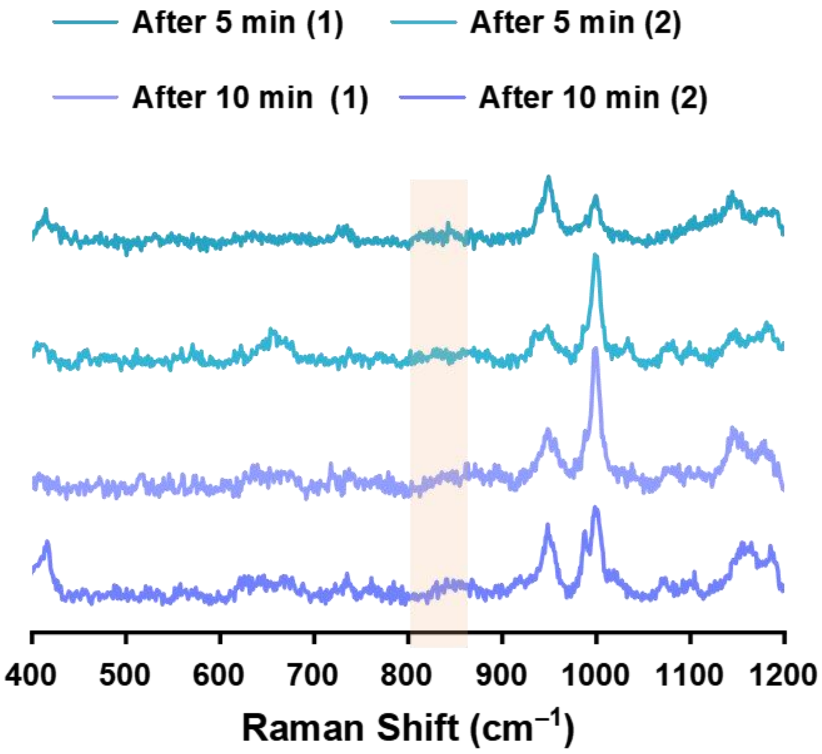


**Supplementary Fig. 21 | SHINERS controls following prolonged exposure of CYP119 to *m*-CPBA.** CYP119 (20 µM) was incubated with *m*-CPBA (40 µM) in 100 mM potassium phosphate buffer (pH 7.0) at 4 °C for 5 or 10 min before SHINERS measurement. Two independently acquired spectra are shown for each incubation time. The shaded region denotes the spectral region containing the ~828 $cm^{-1}$ Fe(IV)=O feature assigned to CYP119 Compound I under the initial reaction conditions. This control tests whether prolonged oxidant exposure reproduces the characteristic ~828 $cm^{-1}$ feature. The absence of an independently resolved corresponding feature argues against a simple assignment of this band to a late oxidant-damaged heme product, although multiple CYP119-derived states may contribute to the overall time-integrated SHINERS spectrum.

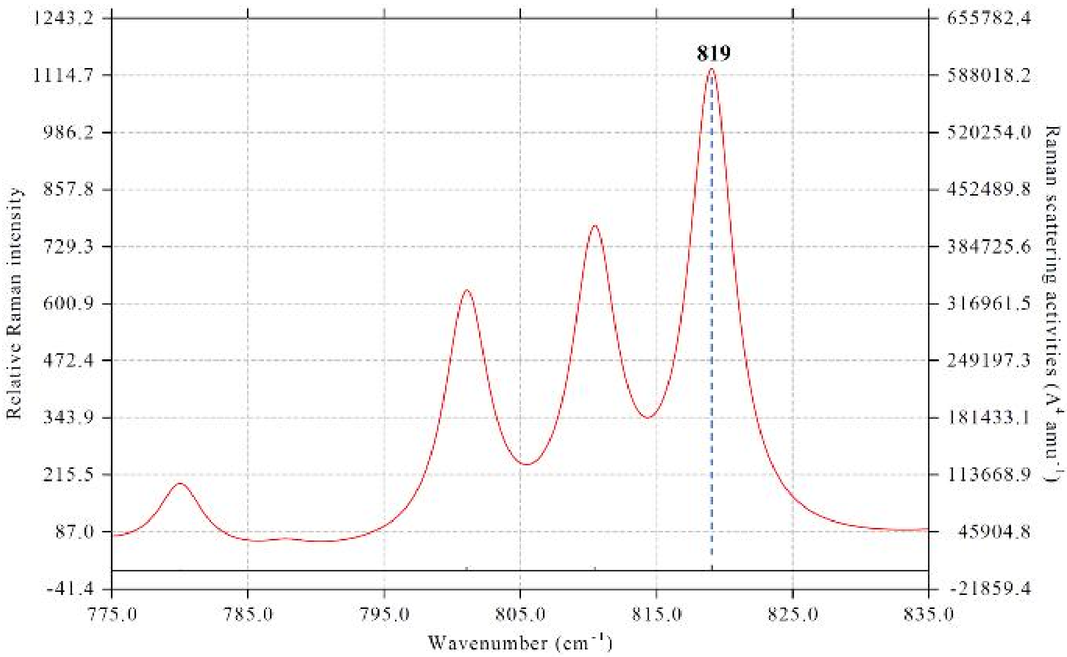


**Supplementary Fig. 22 | Harmonic QM/MM Raman mode calculated for the CYP119 Compound I model.** Calculated Raman activity for the standard-isotope QM/MM-optimized CYP119 Compound I model, showing an Fe–O stretching mode near 819 $cm^{-1}$. No anharmonic correction or empirical frequency-scaling factor was applied. The calculation is used primarily to evaluate Fe–O vibrational mode character and the approximate unscaled frequency. It is not an isotope-substituted normal-mode calculation, and the experimentally observed $^{18}O$-dependent frequency shift is evaluated independently.

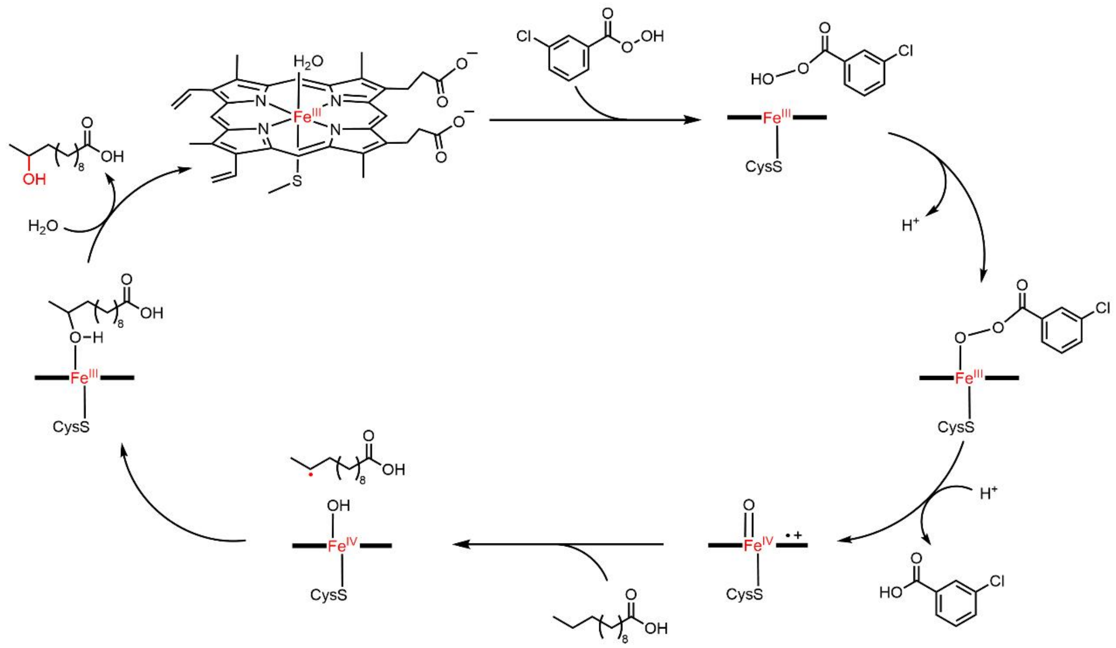


**Supplementary Fig. 23 | Proposed CYP119 catalytic cycle with *m*-CPBA as oxidant.** The scheme summarizes oxidant activation, formation of P450 Compound I, substrate C–H oxidation, product formation, and return to the ferric resting enzyme. The scheme is provided for mechanistic context and does not imply that each intermediate or elementary step is directly resolved by the time-integrated SHINERS measurement.

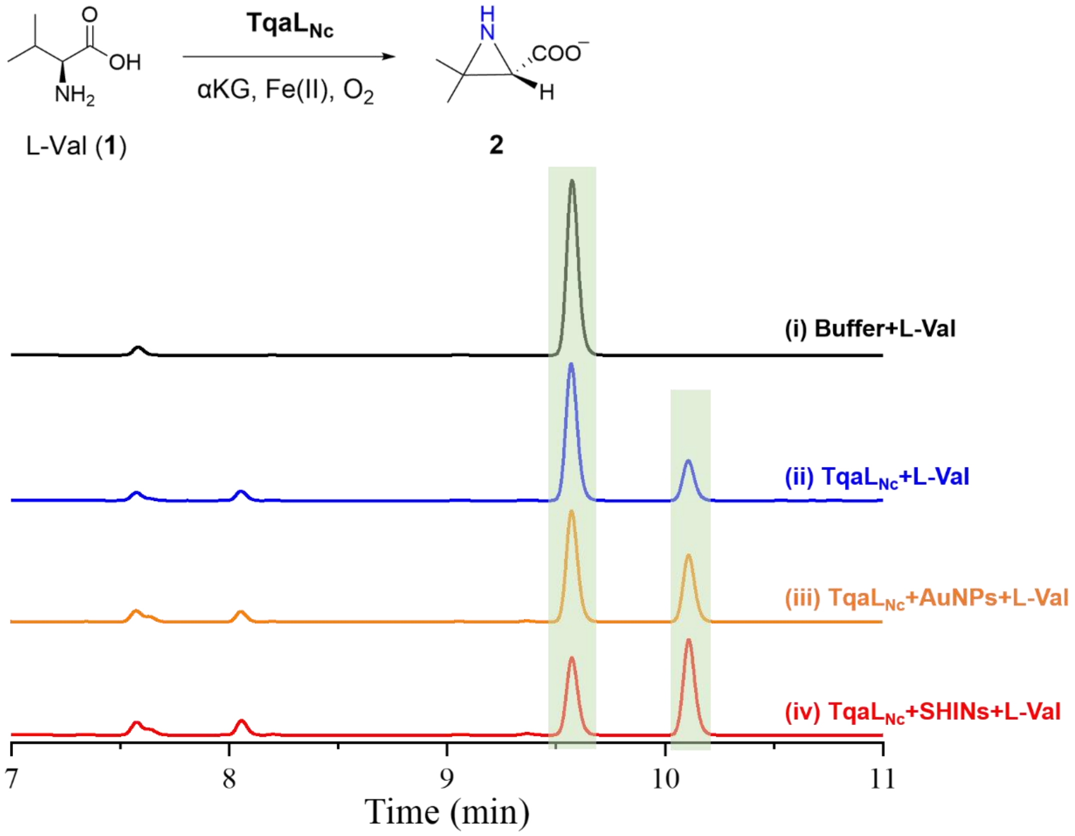


**Supplementary Fig. 24 | Effect of nanoparticles on TqaL$_{Nc}$-catalyzed L-valine aziridination.** HPLC chromatograms for the blank control containing buffer in place of TqaLNc (i), the complete TqaLNc reaction (ii), and complete reactions after pre-incubation with AuNPs (iii) or SHINs (iv). Standard conditions were 20 µM TqaL$_{Nc}$, 1 mM L-Val, 2 mM αKG, and 200 µM $FeSO_4$ in 150 mM Tris-HCl (pH 8.0) with 300 mM NaCl at 20 °C. This measurement serves as a qualitative catalytic-competence control and is not used for quantitative comparison of catalytic rates or for spectroscopic assignment of the Raman features.

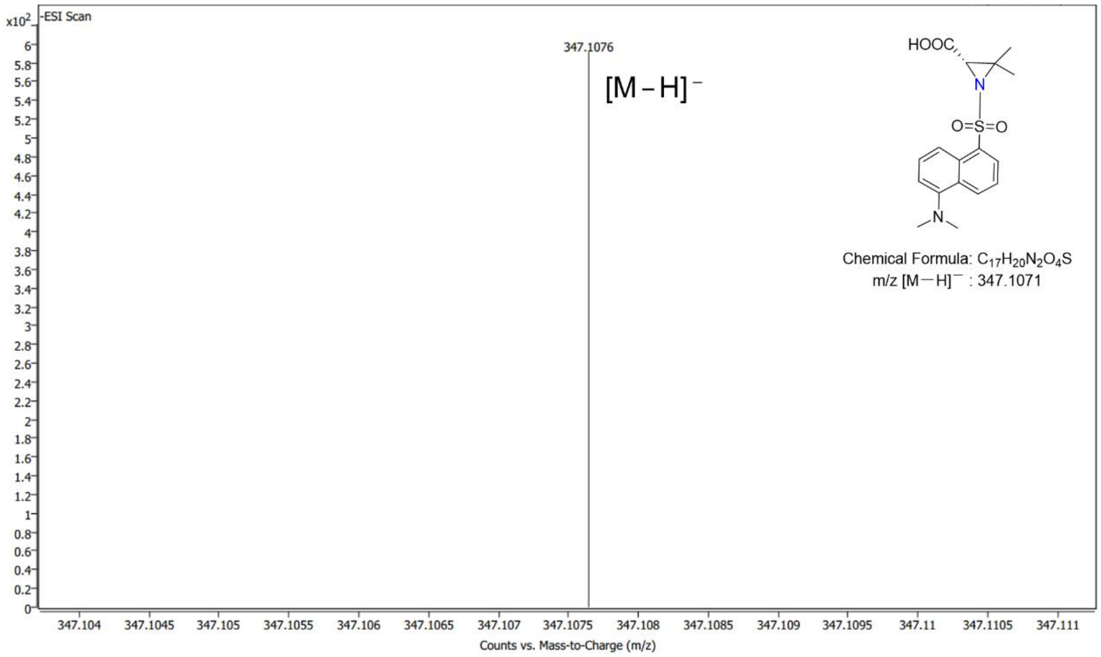


**Supplementary Fig. 25 | LC-MS identification of the TqaL$_{Nc}$ aziridination product.** The TqaL$_{Nc}$ reaction with L-valine was performed at 20 °C for 1 h. The product was derivatized with dansyl chloride and analyzed by LC-MS. The observed ion is consistent with the corresponding dansylated aziridination product.

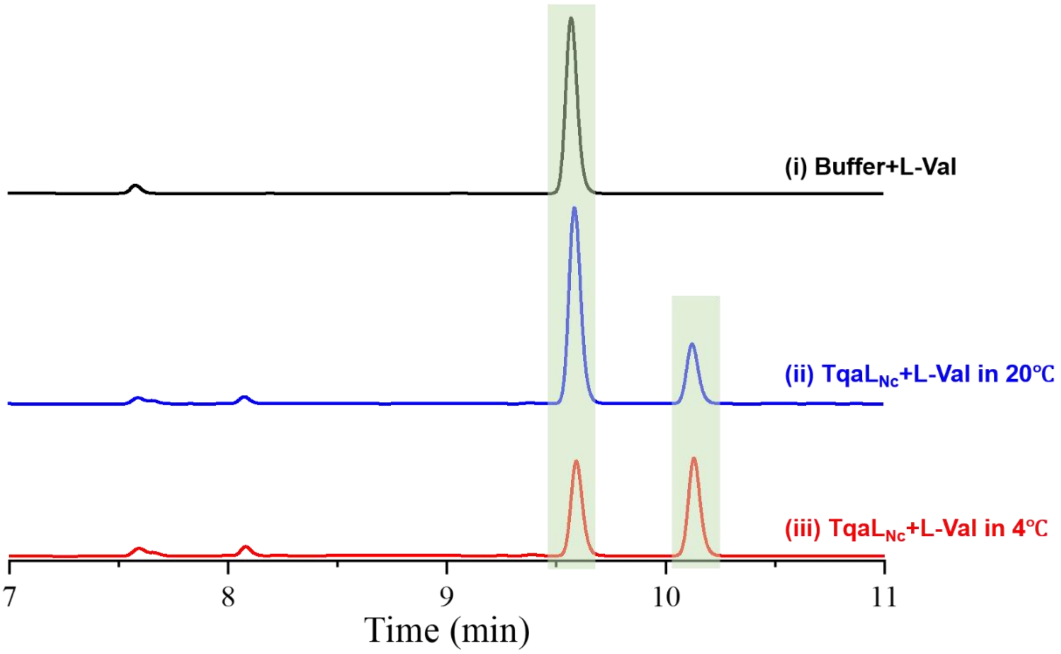


**Supplementary Fig. 26 | Temperature dependence of TqaL$_{Nc}$-catalyzed L-valine aziridination.** HPLC chromatograms for the blank control at 20 °C (i), the complete reaction at 20 °C (ii), and the complete reaction at 4 °C (iii). Reaction mixtures contained 20 µM TqaL$_{Nc}$, 1 mM L-Val, 2 mM αKG, 200 µM $FeSO_4$ in 150 mM Tris-HCl (pH 8.0) containing 300 mM NaCl. This experiment confirms that TqaL$_{Nc}$ product formation remains detectable at 4 °C, the temperature used for the SHINERS measurements, and is not intended to provide temperature-dependent kinetic parameters.

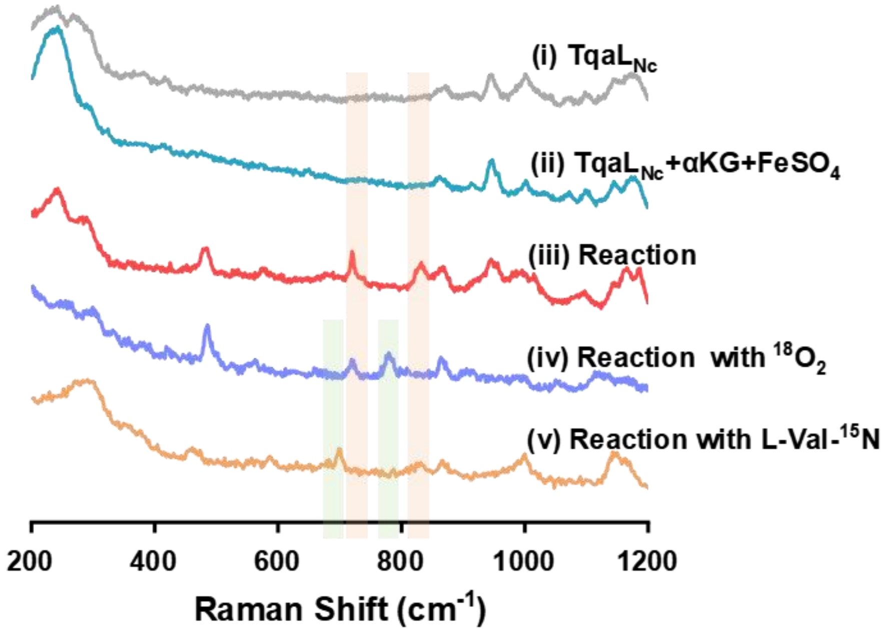


**Supplementary Fig. 27 | Unprocessed raw SHINERS spectra of the TqaL$_{Nc}$ reaction and isotope-substitution experiments.** Spectra shown without baseline correction for i, TqaL$_{Nc}$ alone; ii, the pre-reaction mixture containing TqaL$_{Nc}$, αKG, and $FeSO_4$; iii, the complete reaction with L-Val; iv, the corresponding reaction under an $^{18}O_2$ atmosphere; and v, the reaction with L-Val-$^{15}$N. The shaded spectral regions correspond to the Fe–N- and Fe–O-associated features discussed in the main text. These unprocessed spectra are provided to permit direct assessment of the spectral background and isotope-dependent peak positions independently of baseline correction. Interpretation of the Fe–N- and Fe–O-associated features is based on reproducible, mode-specific isotope responses rather than on changes in overall spectral intensity.

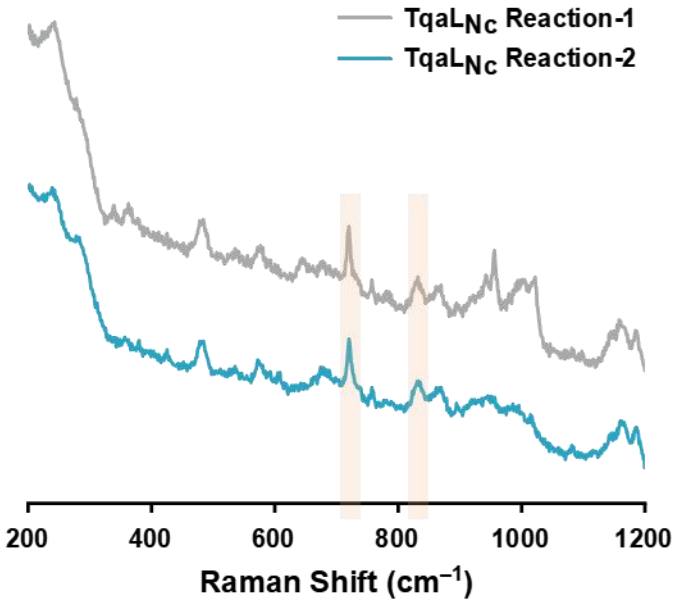


**Supplementary Fig. 28 | Independent unprocessed replicate SHINERS spectra of the TqaL$_{Nc}$ reaction.** Two independently acquired spectra of the complete TqaL$_{Nc}$ reaction containing TqaL$_{Nc}$, αKG, $FeSO_4$, and L-Val are shown without baseline correction. The shaded regions indicate the spectral windows containing the Fe–N- and Fe–O-associated features discussed in the main text. These spectra are included to demonstrate reproducibility of the relevant spectral features in an independent acquisition and are not used to derive statistical uncertainty in the peak positions or isotope shifts.

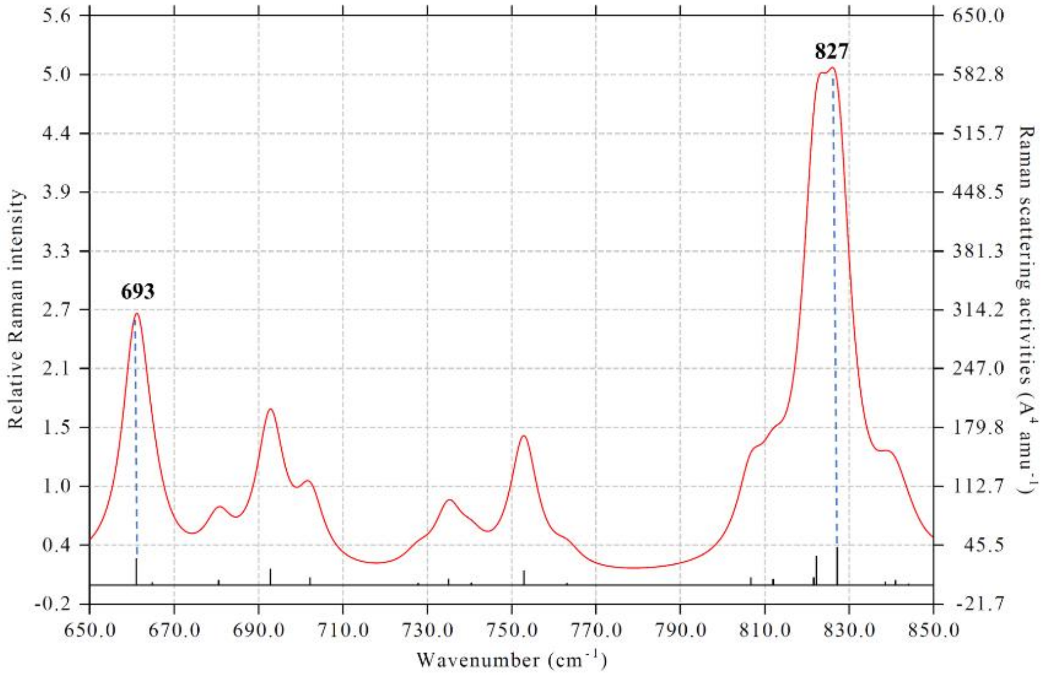


**Supplementary Fig. 29 | Harmonic QM/MM Raman modes calculated for the proposed TqaL$_{Nc}$ (R)HN–Fe(IV)=O state.** Calculated Raman activities for the standard-isotope QM/MM-optimized model, showing modes with Fe–N and Fe–O stretching character near 693 and 827 cm$^{-1}$, respectively. No anharmonic correction or empirical frequency-scaling factor was applied. The calculated modes are used to evaluate Fe–N and Fe–O mode character and approximate unscaled frequencies. They were obtained for the standard-isotope model and are not isotope-substituted normal-mode calculations; the isotope-shift estimates reported in Supplementary Table 1 are independent simplified reduced-mass estimates. Because these vibrations are normal modes of a polyatomic metal–ligand environment, comparison with the reduced-mass estimates is treated as supporting evidence rather than as a pure diatomic-oscillator assignment.

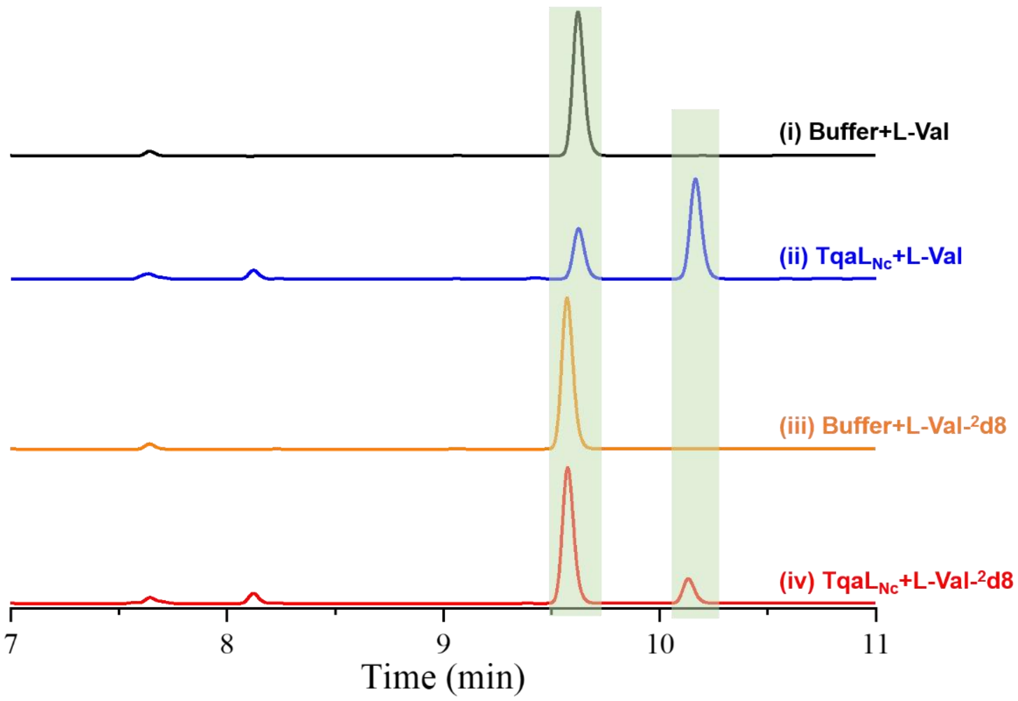


**Supplementary Fig. 30 | Kinetic isotope effect for TqaL$_{Nc}$-catalyzed aziridination.** Representative HPLC analysis of reactions containing 40 µM TqaL$_{Nc}$, 2 mM αKG, 200 µM $FeSO_4$, 1 mM substrate, 150 mM Tris-HCl (pH 8.0), and 300 mM NaCl at 20 °C. i, Blank control with L-Val; ii, complete reaction with L-Val; iii, blank control with L-Val-$^2$d8; and iv, complete reaction with L-Val-$^2$d8. The measured kinetic isotope effect was 4.1. This measurement is included as an independent functional probe of substrate C–H bond cleavage in TqaL$_{Nc}$ catalysis and is not used for assignment of the Fe–N- or Fe–O-associated Raman bands.

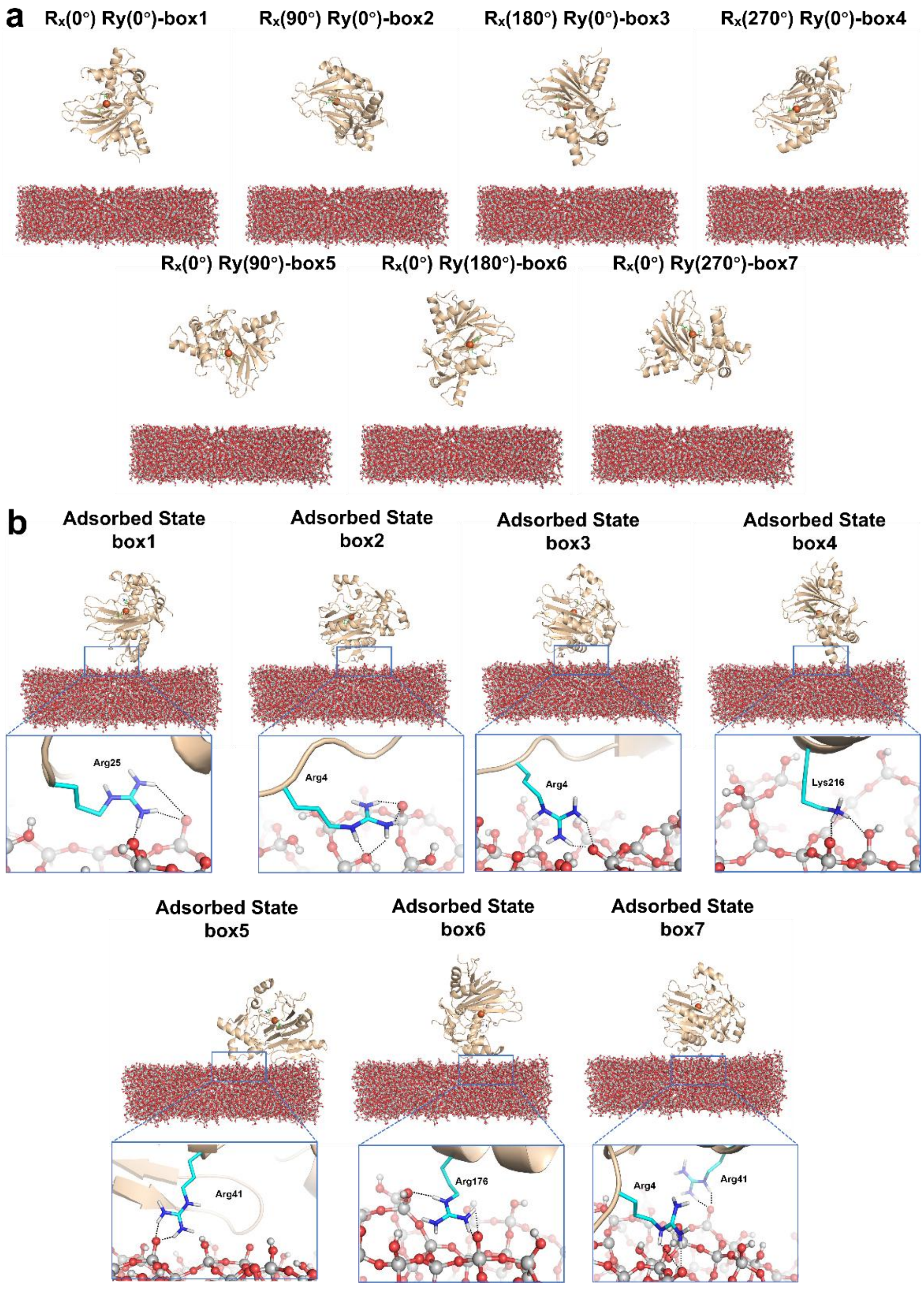


**Supplementary Fig. 31 | MD simulations of TauD adsorption on amorphous silica. a**, Seven initial configurations of TauD above the $SiO_2$ surface, generated by systematic rotations about the x- and y-axes (Rx(0°)Ry(0°) to Rx(0°)Ry(270°)). **b**, Corresponding stable adsorbed structures of TauD from 200 ns MD simulation. Notably, once adsorbed onto the amorphous silica, the enzyme maintains a stable configuration with minor structural fluctuations throughout the subsequent MD simulation.

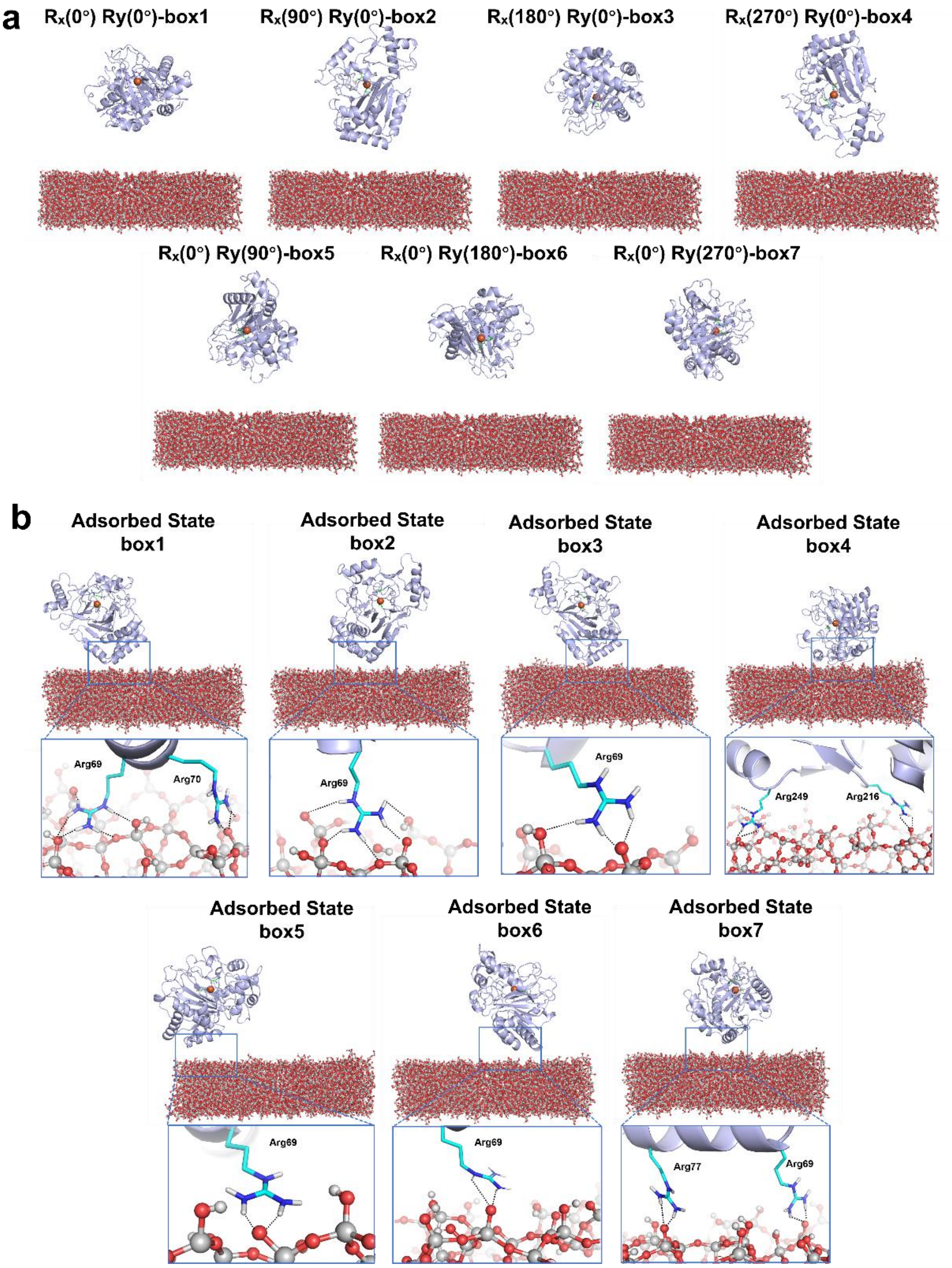


**Supplementary Fig. 32 | MD simulations of VioC adsorption on amorphous silica. a**, Seven initial configurations of VioC above the $SiO_2$ surface, generated by systematic rotations about the x- and y-axes (Rx(0°)Ry(0°) to Rx(0°)Ry(270°)). **b**, Corresponding stable adsorbed structures of VioC from 200 ns MD simulation. Notably, once adsorbed onto the amorphous silica, the enzyme maintains a stable configuration with minor structural fluctuations throughout the subsequent MD simulation.

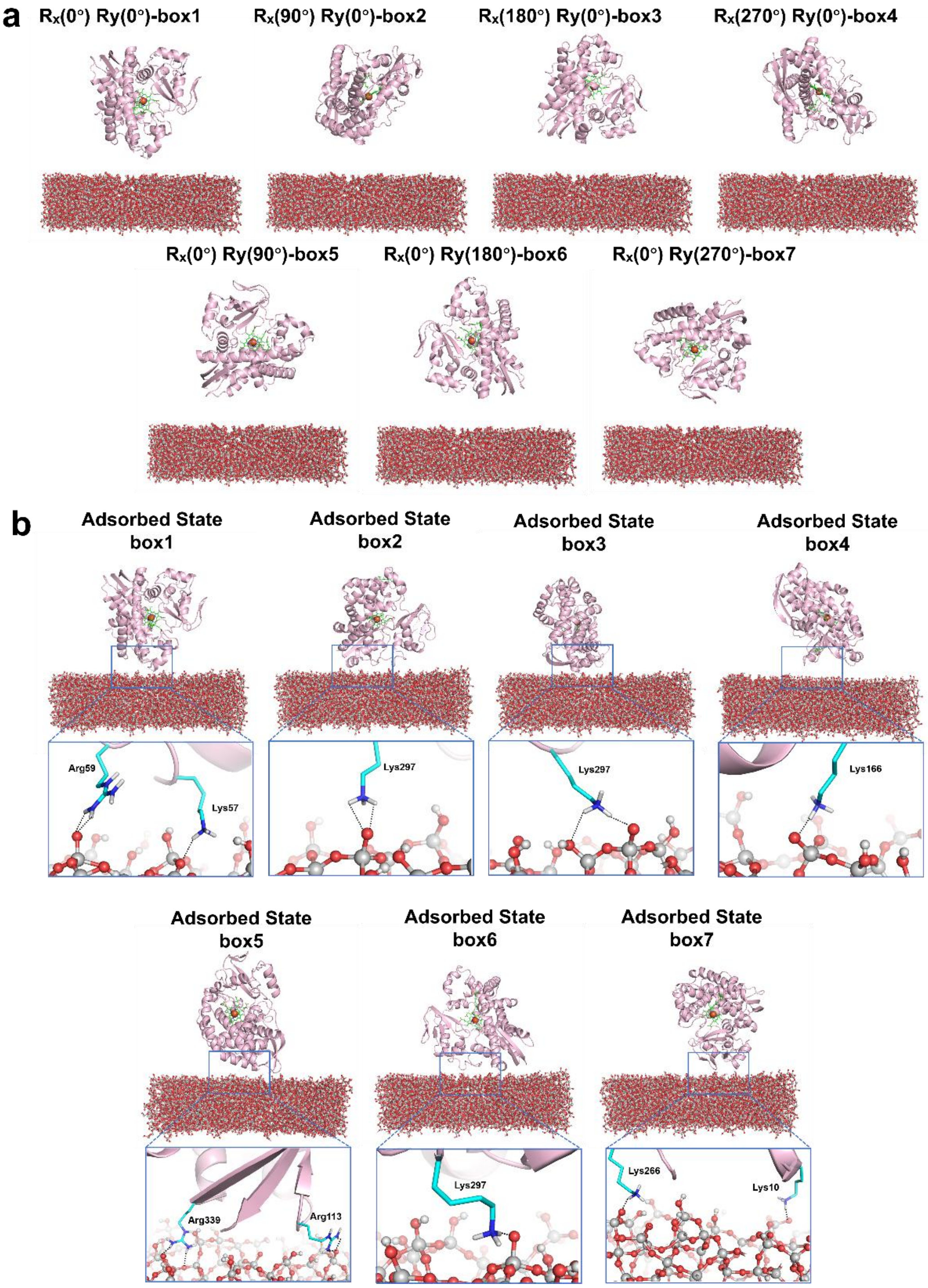


**Supplementary Fig. 33 | MD simulations of CYP119 adsorption on amorphous silica. a**, Seven initial configurations of CYP119 above the $SiO_2$ surface, generated by systematic rotations about the x- and y-axes (Rx(0°)Ry(0°) to Rx(0°)Ry(270°)). **b**, Corresponding stable adsorbed structures of CYP119 from 200 ns MD simulation. Notably, once adsorbed onto the amorphous silica, the enzyme maintains a stable configuration with minor structural fluctuations throughout the subsequent MD simulation.

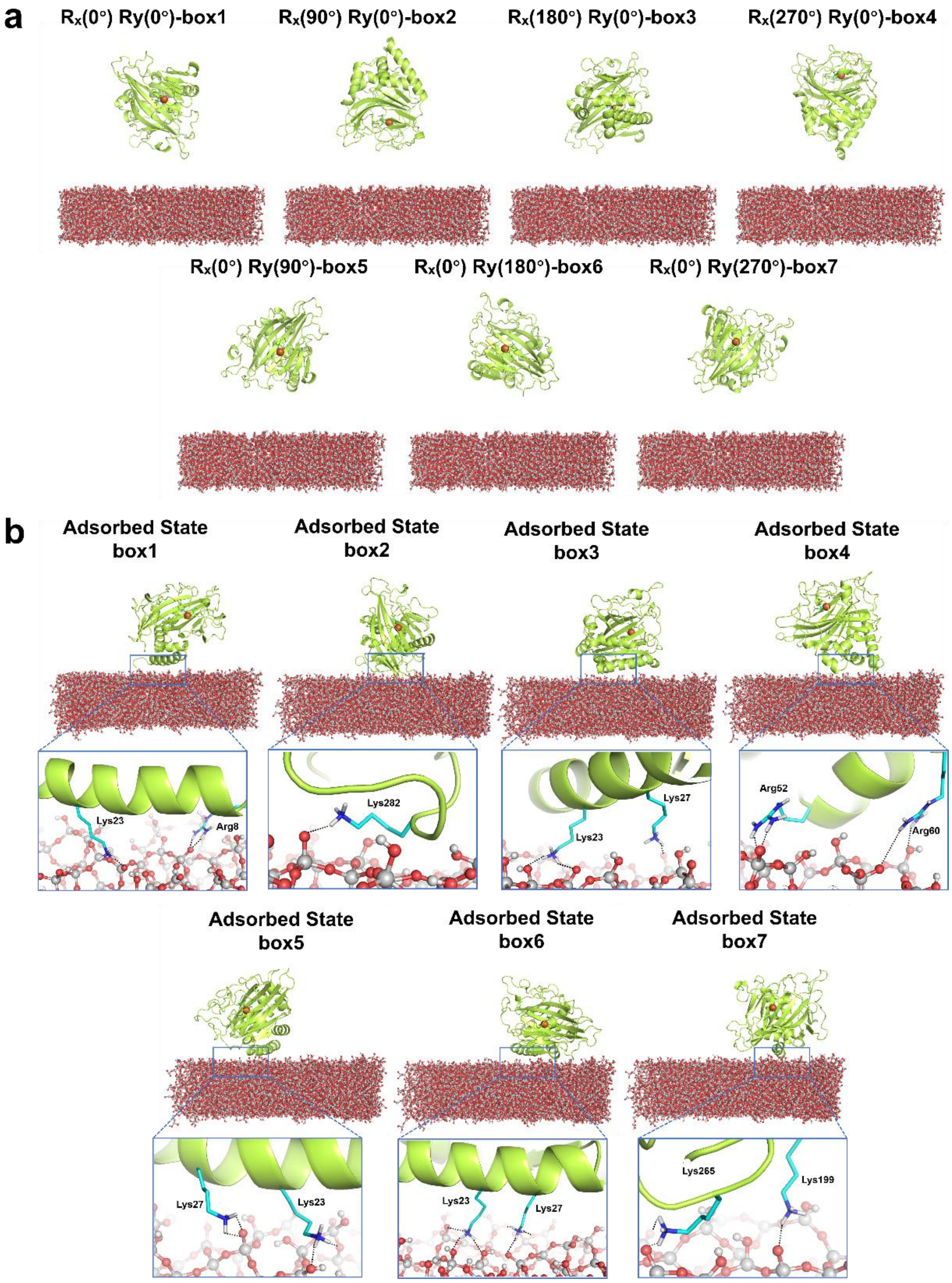


**Supplementary Fig. 34 | MD simulations of TqaL$_{Nc}$ adsorption on amorphous silica.** **a**, Seven initial configurations of TqaL$_{Nc}$ above the $SiO_2$ surface, generated by systematic rotations about the x- and y-axes (Rx(0°)Ry(0°) to Rx(0°)Ry(270°)). **b**, Corresponding stable adsorbed structures of TqaL$_{Nc}$ from 200 ns MD simulation. Notably, once adsorbed onto the amorphous silica, the enzyme maintains a stable configuration with minor structural fluctuations throughout the subsequent MD simulation.

### Supplementary Table

**Supplementary Table 1** Summary of experimental isotope-sensitive Raman features, QM/MM mode character, and spectroscopic interpretation in the Fe-dependent metalloenzyme systems examined in this study.

| System | Experimental mode ($cm^{-1}$) | Isotope-labeled mode ($cm^{-1}$) | Observed shift ($cm^{-1}$) | Simple reduced-mass estimate ($cm^{-1}$) | QM/MM standard-isotope mode | Interpretation |
|---|---|---|---|---|---|---|
| TauD | 582 | 550 ($^{18}O$) | 32 | 26 | — | oxygen-isotope-sensitive feature consistent with Fe(III)–$OO^{-}$ |
| TauD | 822 | 782 ($^{18}O$) | 40 | 36 | ~825, Fe–O stretch | Fe(IV)=O assignment |
| VioC | 816 | 783 ($^{18}O$) | 33 | 36 | Fe(IV)=O structural model | Fe(IV)=O assignment |
| CYP119, no substrate | 828 | 787 ($^{18}O$) | 41 | 36 | ~819, Fe–O stretch | the Fe(IV)=O vibration of CYP119 Compound I |
| CYP119 + lauric acid | 828 | 787 ($^{18}O$) | 41 | 36 | ~819, Fe–O stretch | Fe(IV)=O vibration of CYP119 Compound I under turnover conditions |
| $TqaL_{Nc}$ | 823 | 785 ($^{18}O$) | 38 | 36 | ~827, Fe–O-containing mode | ferryl Fe–O character |
| $TqaL_{Nc}$ | 718 | 701 ($^{15}N$) | 17 | 19 | ~693, Fe–N-containing mode | substrate-N-sensitive mode supporting Fe–N character |

The reduced-mass values are simplified reference estimates and are independent of the QM/MM normal-mode calculations. Because the experimentally observed vibrations are normal modes of polyatomic metal–ligand environments, agreement with a reduced-mass estimate is used only as supporting evidence for isotope-sensitive mode character and is not taken to imply a pure isolated diatomic oscillator. No isotope-substituted QM/MM normal-mode calculations were performed for the present dataset.

### Amino acid sequences

**Sequence 1**: TauD, PDB: 1GQW, GenBank: BAA12841.1

MSERLSITPLGPYIGAQISGADLTRPLSDNQFEQLYHAVLRHQVVFLRDQAITPQQQRALA

QRFGELHIHPVYPHAEGVDEIIVLDTHNDNPPDNDNWHTDVTFIETPPAGAILAAKELPST

GGDTLWTSGIAAYEALSVPFRQLLSGLRAEHDFRKSFPEYKYRKTEEEHQRWREAVAKNP

PLLHPVVRTHPVSGKQALFVNEGFTTRIVDVSEKESEALLSFLFAHITKPEFQVRWRWQPN

DIAIWDNRVTQHYANADYLPQRRIMHRATILGDKPFYRAG

**Sequence 2**: VioC, PDB: 6ALR, GenBank: AAP92493.1

MTESPTTHHGAAPPDSVATPVRPWSEFRLTPAEAAAAAALAARCAQRYDETDGPEFLLDA

PVIAHELPRRLRTFMARARLDAWPHALVVRGNPVDDAALGSTPVHWRTARTPGSRPLSFL

LMLYAGLLGDVFGWATQQDGRVVTDVLPIKGGEHTLVSSSSRQELGWHTEDAFSPYRAD

YVGLLSLRNPDGVATTLAGVPLDDLDERTLDVLFQERFLIRPDDSHLQVNNSTAQQGRVEF

EGIAQAADRPEPVAILTGHRAAPHLRVDGDFSAPAEGDEEAAAALGTLRKLIDASLYELVL

DQGDVAFIDNRRAVHGRRAFQPRYDGRDRWLKRINITRDLHRSRKAWAGDSRVLGQR

**Sequence 3**: CYP119, PDB: 1IO7, GenBank: AAB03278

MYDWFSEMRKKDPVYYDGNIWQVFSYRYTKEVLNNFSKFSSDLTGYHERLEDLRNGKIR

FDIPTRYTMLTSDPPLHDELRSMSADIFSPQKLQTLETFIRETTRSLLDSIDPREDDIVKKLA

VPLPIIVISKILGLPIEDKEKFKEWSDLVAFRLGKPGEIFELGKKYLELIGYVKDHLNSGTEV

VSRVVNSNLSDIEKLGYIILLLIAGNETTTNLISNSVIDFTRFNLWQRIREENLYLKAIEEALR

YSPPVMRTVRKTKERVKLGDQTIEEGEYVRVWIASANRDEEVFHDGEKFIPDRNPNPHLS

FGSGIHLCLGAPLARLEARIAIEEFSKRFRHIEILDTEKVPNEVLNGYKRLVVRLKSNE

**Sequence 4**: $TqaL_{Nc}$, PDB: 7EEH, GenBank: CAD21320.1

RLDVTKRVFNPSYYTAIAEIMKLRSKYITNRSIFVEGSDMVPLLLGLGATRADLDALQRVS

NNLYSDPTLPFRRSRNGRFCFDFSTRSVRRLEFQPFALSVEEDFKRHDSGQIRVFDEVQDEL

QLNTAFQALLVFKGMICHGVQTTHRPRLDYSSDKWVCTLFNLRTVTTPSILGEPALEGVH

TDGVDHTMTTYLGSKNMDLAANSAVTFMHDMNEETGAKYTEIKPQNLRSRVQHRHFLD

TLLLVDTENKHSLSPVLPLDETKEATRDMLIFFTRRPVKKGDGGHISENIDSFRPHEELPME

VPLFLPGA
